# Neuromodulation of risk preferences encoded in human orbitofrontal cortex activity

**DOI:** 10.1101/2025.05.06.650317

**Authors:** Alexandra Skular, Lu Jin, Jacqueline A. Overton, Ignacio Saez

## Abstract

Human behavior presents a natural range of variation beyond which extreme behavior appears, often as a hallmark of psychiatric conditions. for example, risk preferences, which naturally range from risk-seeking to risk-averse, are characteristically affected in multiple pathologies, e.g. increased risk-taking in gambling disorders. Despite its basic and translational importance, how underlying brain activity reflects and determines behavioral variation is mostly unknown, in part due to the difficulty of directly examining neural activity in decision-relevant brain areas from cohorts of subjects with varying preferences. To address this, we combined human intracranial recordings (n=15 patients) from the orbitofrontal cortex (OFC), a key brain region in economic decision-making, with risky decision-making tasks and computational modeling. We observed significant behavioral variation in risk preferences, which was strongly correlated with low-frequency (delta-theta) oscillatory coherence in OFC. Low-frequency oscillatory phase was coupled to high-frequency activity amplitude, providing a potential neurophysiological mechanism for integrating preferences with trial-by-trial risk computations. Finally, targeted OFC electrical stimulation caused significantly decreased risk-taking behavior and faster reaction times without impairing overall decision-making. These results show a neural correlate for inter-individual variation in decision-making preferences and provide proof-of-principle evidence for modulation of risk-taking behavior through OFC-targeted neurostimulation, paving the way for developing neuromodulatory interventions for pathologies characterized by heightened risk-seeking, such as addiction and gambling disorders.

**Significance statement:** We leverage invasive electrophysiological recordings from human orbitofrontal cortex to demonstrate that low-frequency oscillatory coherence reflects stable inter-individual economic preferences, with targeted neurostimulation causing risk averse behavior. These results provide a neurobiological substrate for interindividual differences in economic decision-making and provide a potential therapeutical avenue for brain disorders characterized by risky behavior.

## Introduction

Aberrant decision-making behavior is a hallmark symptom of many psychiatric conditions, including depression, gambling, addiction and obsessive-compulsive disorders (1, 2). Specific aspects of decision-making are affected across multiple diagnoses, suggesting they may share a common underlying neurobiological mechanism. For example, the amount of risk individuals are willing to tolerate for a chance to obtain an uncertain reward (3, 4), extends beyond a normal range in certain pathologies: depression is often characterized by risk-averse behavior (5), while gambling addiction is associated with risk-seeking behavior(6). Therefore, a mechanistic depiction of the neural basis of risk preferences would be valuable, both as a normative description of natural variation (1) and as a biomarker for psychiatric pathologies in which this behavior is affected (5, 6). In addition, it could open the door to development of novel invasive neurostimulation techniques to treat psychiatric disorders characterized by extreme behavior, by focusing on approaches capable of correcting behavioral deficits (7).

Despite its importance, the underlying neural basis of behavioral preferences, including risk attitudes, are not well understood. Multiple converging lines of evidence indicate that the human orbitofrontal cortex (OFC) plays a central role in decision-making under uncertainty and economic behavior (8, 9). Both animal and human experiments demonstrate that OFC activation encodes information related to risk in choices under uncertainty as measured by fMRI BOLD (10), single unit spiking (11), and high-frequency gamma activity (12). In contrast to trial-by-trial risk computations relevant to ongoing decisions, individual risk preferences are a stable, temporally sustained trait (3, 13), making fast, trial-level modulations of neural activity poor candidates for representing risk preferences, suggesting a different neural substrate may be at play. Low frequency oscillations are slow, longer-range neural features capable of establishing functional communication through coherent oscillations(14, 15) and modulating high frequency activity through cross frequency coupling (CFC) (16). Low frequency oscillatory dynamics are implicated in a variety of cognitive processes (17, 18), with abundant evidence implicating activity in the theta-band range (θ, 4-8Hz) in decision-making (19–21). Prior evidence suggests that inter-individual differences in cognition and behavior may be functionally expressed by differences in oscillatory patterns in a variety of contexts, including motivation (22), working memory (23), conflict processing (18), and moral decision-making (24). Therefore, low frequency oscillations are a well suited neurophysiological substrate for the neural representation of risk preferences. In addition, intracranial recordings from human populations present a unique opportunity to examine the neural basis of inter-individual behavioral variation. Non-human primate (NHP) studies, though instrumental in demonstrating a role of OFC in reward-related behavior (8, 9), suffer from practical limitations in numbers of subjects (typically n=2-3) restricting the study of inter-individual variation. Conversely, non-invasive human studies (fMRI, EEG) are limited in temporal resolution, neurophysiological detail and access to deep brain structures, such as OFC, that are likely to be central for encoding behavioral preferences.

Here, we leveraged the unique qualities of invasive neurosurgical interventions (intracranial electroencephalography, iEEG) in human patients, which provide electrophysiological data (local field potentials, LFPs) with high anatomical specificity, temporal resolution, and neurophysiological detail (25), to identify the precise oscillatory dynamics involved in human risk-taking behaviors. By combining iEEG with behavioral performance in a risky decision-making task (12, 26), we investigated the role of OFC low frequency oscillations in representing individual risk preferences. We found that inter-individual variation in intra-OFC coherence in the delta-theta (δ-θ, 1-8Hz) frequency range wa s strongly correlated with inter-individual risk preferences, with individuals with higher δ-θ coherence exhibiting riskier behavior. The phase of δ-θ oscillations was consistently coupled to the amplitude of broadband gamma activity (γ-*H*γ, 30-200Hz), mechanistically linking representations of stable risk preferences in low frequency oscillatory phase and trial-by-trial risk in high frequency encoding. Finally, using OFC-targeted electrical stimulation, we modulated patients’ risk-taking behavior, causing risk-averse behavior. These findings suggest a mechanism for representation of risk preferences in the human OFC and demonstrate the ability to selectively alter risk preference through invasive OFC stimulation (Fig. 5). Overall, we provide a proof-of-principle framework for the efficacy of neuromodulation in modulating human choice behavior, highlighting the therapeutic potential for invasive neuromodulation interventions to treat disorders characterized by aberrant risk preferences.

## Results

### Neurosurgical patients show variation in risk preferences

We recorded electrophysiological LFP activity from epilepsy patients (*n*=15, *n*=5 female; age µ=35.2±8.30) undergoing iEEG monitoring (stereotactic EEG, sEEG *n*=7/15 or electrocorticography, ECoG *n*=8/15) while they played a risky decision-making game (Fig. 1a). We selected patients with OFC coverage (Fig.1b). After confirmation of anatomical location and electrophysiological quality control (see Methods), our final dataset was comprised of *n*=130 OFC electrodes (µ=8.67 electrodes/patient; see Extended Data Table 1).

**Figure 1.**
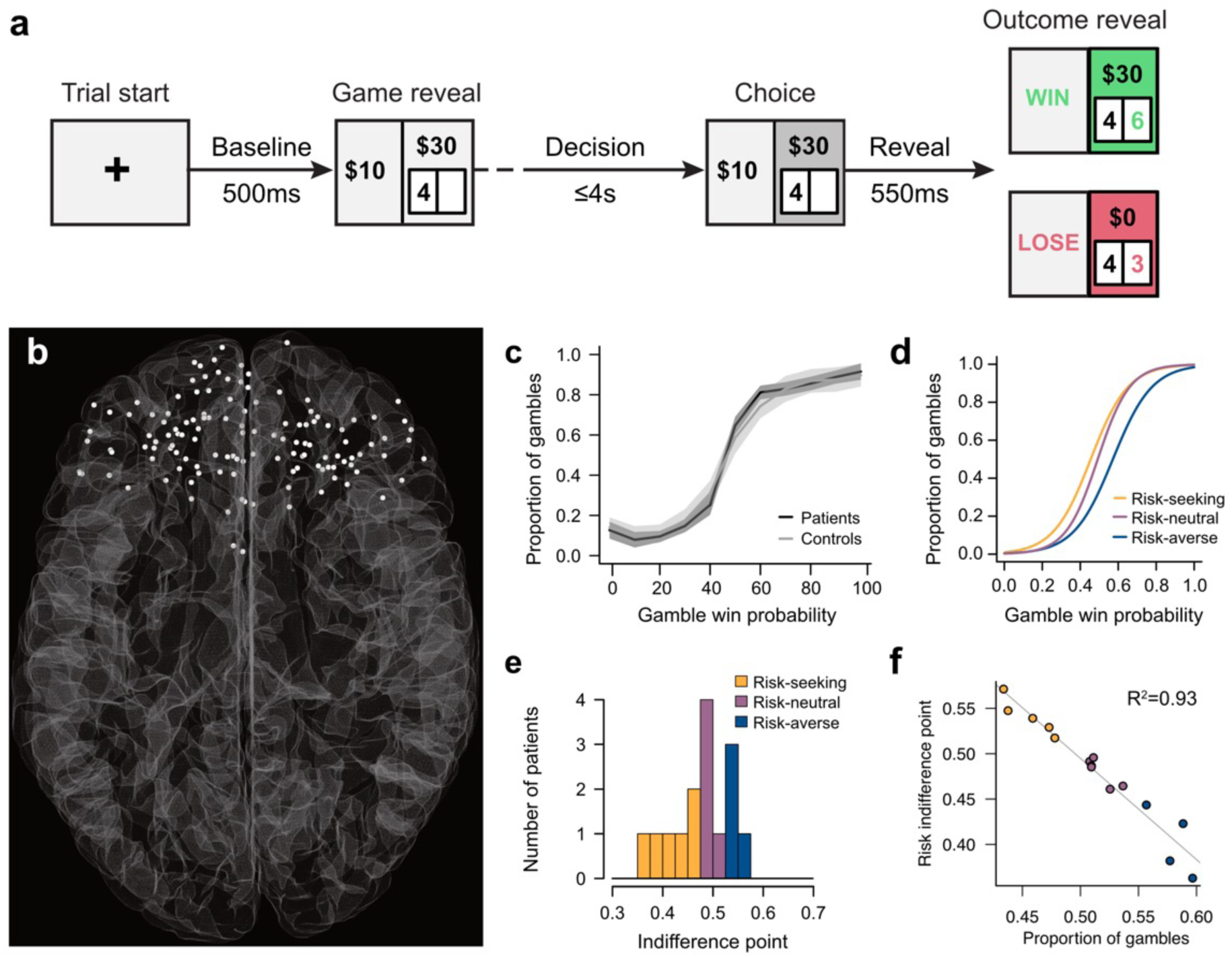
Experimental methods: iEEG recording and characterization of inter-individual risk attitudes during gambling task. **a**, Behavioral task design. Patients played a gambling task in which they chose between a safe prize of $10 and a risky bet for a higher prize (e.g. $30). Players played a total of 200 rounds, in which the win probability varied between 0-100% in 10% increments. **b**, Anatomical reconstruction showing the location of all intracranial iEEG electrodes located in orbitofrontal cortex (OFC) included in our sample (*n*=130, *n*=15 patients). **c**, Average patient behavior (dark grey line) compared to healthy controls (light grey line). Patients chose to gamble more often for trials in which the win probability was higher (*P*<0.001; random effects logit analysis; shaded area = SEM). Choices were comparable to those of healthy participants (all *P*>0.2). **d**, Individual patient plots reveal inter-subject variability in risk preference, showing example patients’ risk-seeking (yellow), risk-neutral (magenta) or risk-averse (blue) behavior. **e**, Distribution of patients’ risk indifference points, defined as the win probability at which they’re equally likely to choose the safe bet or the risky option. Patients showed risk-seeking, risk-neutral and risk-averse behavior. **f**, Increasingly risk-seeking behavior was associated with a greater proportion of gambles (*R^2^*=0.93, *P*<10E^-8^).

Patients played a gambling task in which they chose between a safe bet and a risky gamble (Fig.1a). Win probability varied parametrically from trial to trial between 0% (sure loss) and 100% (sure win) in 10% increments (*n*=200 rounds). Patients chose to gamble more often when the expected value of the gamble was higher (*P*<0.001, random effects logit analysis, Fig.1c), indicating that they understood and could play the game. We compared behavior in our epileptic cohort to a healthy control cohort (*n*=20) and found no significant difference in the proportion of gambles for all win probabilities (all *P*>0.2; t-test; Fig.1c), indicating our patients behaved in a manner comparable to healthy controls. To measure risk preferences across patients, we calculated each patients’ risk indifference point, a model-free estimate of risk preference defined as the win probability for which the player is equally likely to select the gamble or the safe bet. Risk-neutrality is indicated by a risk indifference point of 0.5, at which players are equally likely to choose the risky gamble or the safe bet in 50% win probability gambles. In contrast, risk indifference points below/above 0.5 indicate risk-seeking/risk-averse behavior, respectively. In addition, we also measured risk preferences using a model-based strategy, using a prospect theory-derived model (27) that captures individual risk preferences as a risk aversion parameter (see Methods). There was a high correlation between our model-free and model-based risk preference metrics (*R^2^*=0.975, *P*<0.001), indicating they are similarly capable of capturing inter-individual variation in risk preferences (Extended Data Fig.1a,b). To obtain a sense of optimal play, we generated task behavior from artificial agents of varying risk preferences (see Methods, Extended Data Fig.1c,d). Simulated agents with a risk indifference point of 0.43 obtained maximum profits in the task (Extended Data Fig.1d). On average, iEEG patients’ risk preferences were close to optimal (risk indifference point µ=47.9%±5.29%; max=57.12%, min=36.28%; Fig.1d).

Next, we examined variation in play across patients. For descriptive purposes, we classified subjects’ risk preference as risk-seeking (risk indifference point ≤0.475), risk-neutral (risk indifference point between 0.475 and 0.525) or risk-averse (risk indifference point ≥0.525). Most patients were risk-seeking (*n*=6/15) or risk-neutral (*n*=6/15), with a lower proportion of patients displaying risk-aversion (*n*=3/15; Fig.1e). To confirm risk indifference points accurately reflected choice behavior, we tested whether there was a correlation between a patient’s propensity to gamble and their risk indifference point. As expected, we found that patients with higher risk indifference points (risk-seeking) had significantly higher proportions of gamble choices during play (Fig.1f; *R^2^*=0.937, *P*<10^-8^).

### Intra-OFC **δ**-**θ** coherence correlates with risk preferences

Based on prior evidence relating low frequency oscillations to decision-making behavior (19) and inter-individual differences (18, 24), we hypothesized that OFC low-frequency oscillations may represent a stable neural substrate for variation in risk preferences. We focused on coherence, a metric hypothesized to reflect functional connectivity(14, 15), and in the delta-theta (δ-θ, 1-8Hz) frequency range, previously implicated in decision-making (19–21). To obtain an OFC-wide measure of coherence, we calculated the average δ-θ coherence separately for each patient. Specifically, we computed the mean pairwise δ-θ coherence between all OFC electrodes within-subject (*n*=130 electrodes across *n*=15 subjects, see Extended Data Table 1) during the decision epoch (-1 to 0 seconds prior to choice button press; Fig.1a), and found considerable variation in OFC-wide δ-θ coherence across patients (µ=0.29±0.087; Extended Data Fig.2).

Next, we related δ-θ coherence to individual risk preferences, measured by the risk indifference point (Fig.2a). We found a negative correlation between mean OFC δ-θ coherence and risk indifference point, with risk-seeking patients showing greater intra-OFC δ-θ coherence (Fig.2a; *R^2^*=0.61, *P*=0.0006, *n*=15). This relationship was also present, albeit slightly weaker, between δ-θ coherence and the prospect theory-derived risk preference parameter (*R^2^*=0.49, *P*=0.002, see Methods). To examine whether this relationship extended beyond the decision epoch, we carried out the same analysis in the baseline epoch (-0.5 to 0 seconds before stimulus presentation). We found that this relationship was also present, although attenuated, during baseline (Fig.2b; µ=0.22±0.058, *R^2^*=0.44, *P*=0.007, *n*=15), indicating that the relationship between δ-θ coherence and risk preferences was stable beyond the decision period.

To probe the specificity of this relationship, we carried out several control analyses. First, to ensure that we correctly estimated slow δ oscillations, we repeated our analysis with a longer decision epoch (3s prior to button press) that encompassed a minimum of three full cycles for the slowest δ oscillation (1Hz) in our target frequency range (δ-θ, 1-8Hz). Using this longer window, the correlation between δ-θ coherence and risk indifference point was maintained (*R^2^*=0.45, *P*<0.01, Extended Data Fig.3). Because δ and θ are often considered separately, we tested the association between coherence and risk indifference point in each frequency band separately. We found that the negative correlation was still present, albeit slightly weaker, in both frequency bands (*R^2^*=0.59 and *R^2^*=0.46 for δ and θ, respectively; both *P*>0.01; Extended Data Fig.4a-b,e-f). We examined whether intra-OFC representation of risk preference was specific to the δ-θ frequency band by extending our analysis to other low-frequency bands, namely α (9-12Hz) and β (13-30Hz). We found that the relationship between coherence and risk preferences in α or β OFC coherence was not significant or very weak (Extended Data Extended Data Fig.4d-c,g-h), indicating that this relationship is mainly present in the δ-θ frequency band. Next, we investigated whether this relationship was specific to low frequency oscillatory coherence, rather than mere oscillatory power related to the 1/f profile of neural activity. We observed no significant correlation between risk preferences and average intra-OFC δ-θ power (*P*>0.3, Extended Data Fig.5). Finally, we investigated whether risk preference representation was specific to the OFC or reflected brain-wide oscillatory patterns by performing a similar analysis in hippocampal contacts. We observed no correlation between hippocampal δ-θ coherence and risk preference (Extended Data Fig.6), suggesting that risk preference is not widely represented across brain areas.

### **δ**-**θ** phase modulates high frequency **γ**-*H***γ** amplitude

Our observation that risk preferences are associated with intra-OFC δ-θ coherence raises the question of how this neural representation is integrated with trial-specific information about immediately available choices. Our previous study determined that OFC encodes trial-by-trial reward computations, including risk and win probability, through modulations in high frequency broadband gamma (γ-*H*γ, 30-200Hz) power. Together with our δ-θ coherence results, this suggests that integration may be achieved through functional interactions between frequency bands. Phase-amplitude cross frequency coupling (CFC), whereby the phase of low frequency oscillations modulates the amplitude of fast, high frequency activity, is commonly observed in human cortex (16, 28, 29) and could play this role. Therefore, we hypothesized that phase-amplitude CFC is involved in integrating risk preference (reflected in δ-θ oscillatory phase) and reward computations (encoded by high frequency γ-*H*γ activity) in the OFC.

To probe possible phase-amplitude relationships across frequencies, we measured CFC between a range of low frequency phases (2-20Hz) and high frequency amplitudes (5-200Hz). For every low frequency phase (*lf_p_*) and high frequency amplitude (*hf_a_*) combination (*n*=760), we quantified phase-amplitude coupling as the mean phase-locking value (PLV) between each *lf_p_*:*hf_a_*pair during the decision epoch (-1 to 0 seconds prior to choice, Fig.1b) within every OFC electrode (*n*=130) for all subjects (*n*=15; see Methods). We used a surrogate distribution approach to evaluate the significance of the observed PLV for each *lf_p_*:*hf_a_* through non-parametric statistical testing (see Methods). Group-level statistical tests identified electrodes with significant CFC patterns if there was at least one significant cluster consisting of any *lf_p_*:*hf_a_* pairs, after surrogate normalization (see Methods).

We found robust patterns of significant phase-amplitude coupling between low frequency phases and high frequency amplitudes across electrodes. A representative electrode with strong δ-θ:γ-*H*γ phase-amplitude coupling is shown in Fig.3. This pattern was common, with 36/130 OFC electrodes (27.70%) showing significant CFC (see Methods). To estimate consistent patterns and identify the main frequencies involved in CFC, we aggregated PLV_z_ results across electrodes showing significant CFC in any *lf_p_*:*hf_a_* combination (Fig.3b). In this group analysis, CFC was strongest between δ-θ *lf_p_* phases and γ-*H*γ *hf_a_* amplitudes (Fig.3b), a pattern that is absent in electrodes without significant CFC (Extended Data Fig.7a).

To further determine which phase and amplitude frequencies had the strongest cross-frequency interactions, we computed the marginal average PLV_z_ distribution across all *hf_a_* amplitude frequencies and for all *lf_p_* phase frequencies. We found that CFC was greatest in two pairs of peaks in the high and low-frequencies: low gamma (γ, 35Hz) and high gamma (*H*γ, 90Hz) in *hf_a_* frequencies (5-200Hz; Fig.3c), and delta (δ, 3Hz) and theta (θ, 5Hz) in *lf_p_*frequencies (Fig.3d). Therefore, cross frequency phase modulation of high frequency amplitude was strongest in δ-θ, the same frequency range which showed the strongest association with individual risk preference (Fig.2).

**Figure 2.**
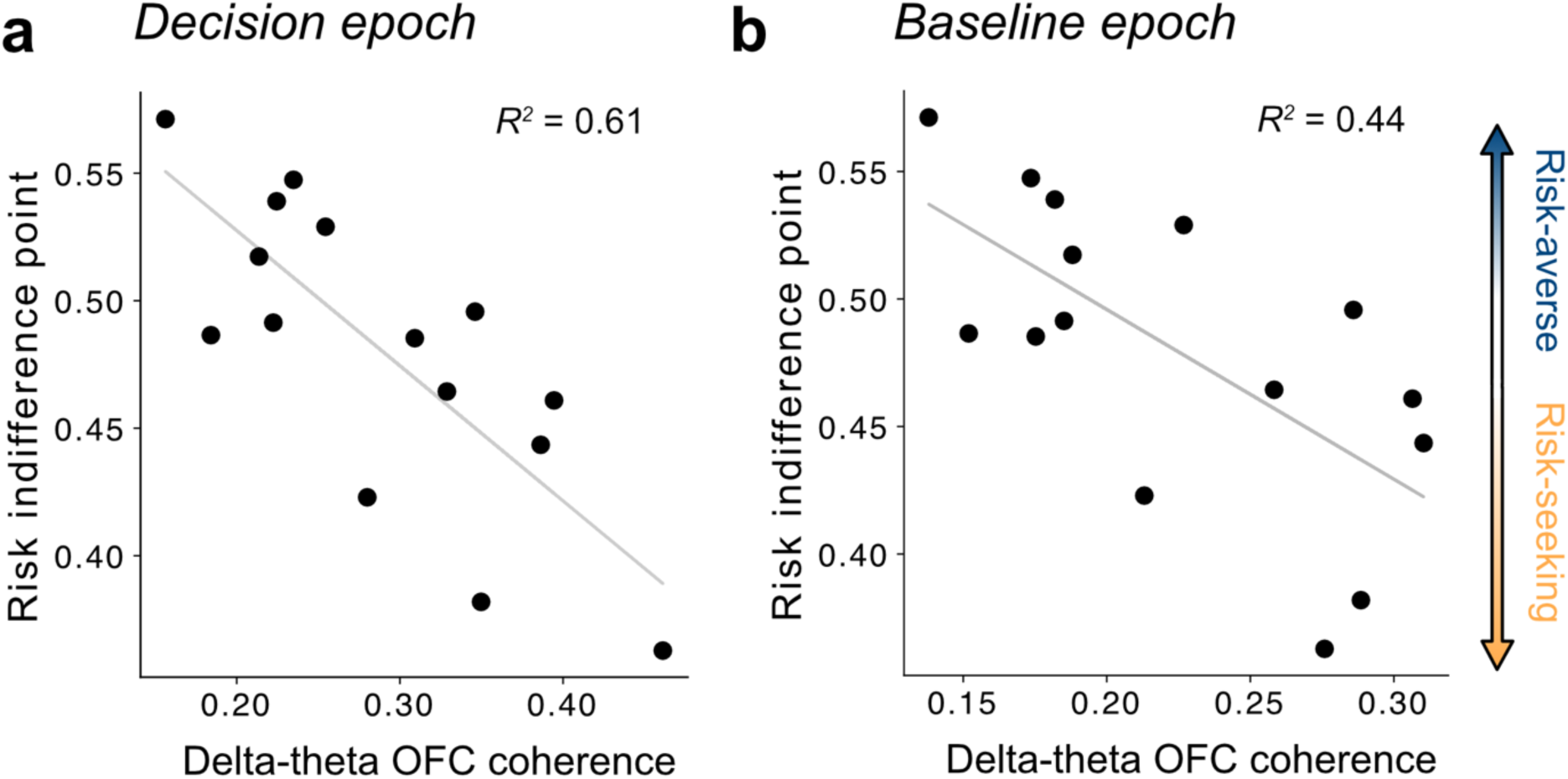
Representation of individual risk preferences in intra-OFC delta-theta coherence. a-b, We calculated the average delta-theta (δ-θ, 1-8Hz) intra-OFC coherence for each patient as the mean pairwise δ-θ coherence between all OFC electrodes in each patient. Each point represents a single patient (*n*=15). We observed a significant negative correlation between subjects’ average OFC δ-θ coherence and their risk indifference point, with patients with higher δ-θ coherence showing riskier behavior. This relationship was present during the 1s pre-choice deliberation epoch (**a**) (*R^2^*=0.61,*P*=0.0006) as well as the 0.5s pre-trial baseline epoch (**b**) (*R^2^*=0.44,*P*=0.007).

**Figure 3.**
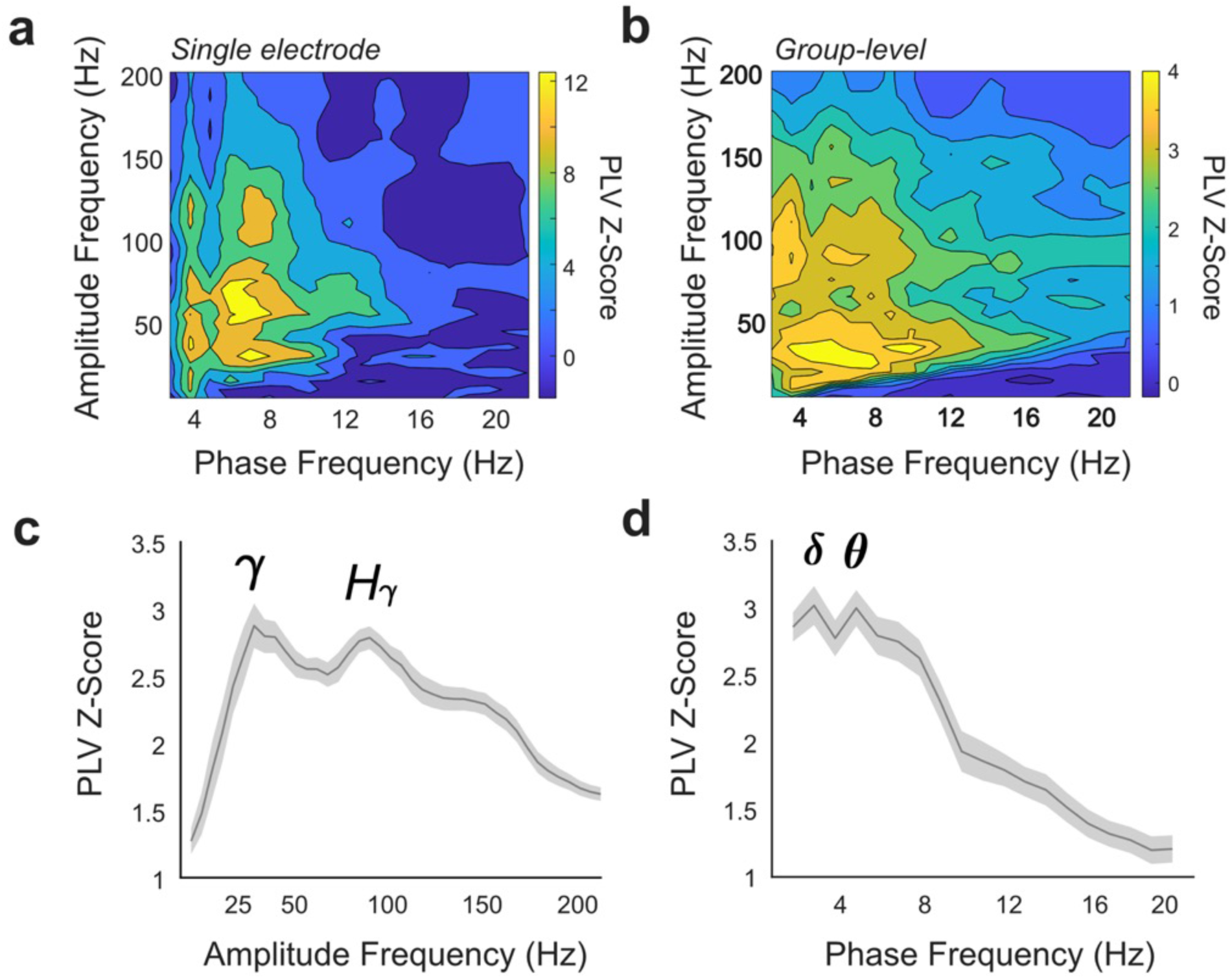
Phase-amplitude cross frequency coupling between low frequency delta-theta phase and high frequency broadband gamma amplitude. Cross frequency coupling was calculated as the phase-locking value (PLV) between all combinations of low frequency phases (*lf_p_*, 2-20Hz) and high frequency amplitudes (*hf_a_*, 5-200Hz) during the decision epoch across trials for every OFC electrode (*n*=130) across subjects (*n*=15). PLVs were normalized (PLV_z_) using their null surrogate distribution (*n*=1000, see Methods). **a**, Example comodulogram from a representative electrode, representing PLV_z_ for each *lf_p_*:*hf_a_* pair (*n*=760). **b**, Group-level comodulogram of average PLV_z_ from electrodes with significant CFC across subjects (*n*=36/130 electrodes from 10/15 subjects). An electrode was classified as significant if there was at least one significant comodulogram cluster (see Methods). **c**, Distribution of average PLV_z_ for all high frequency amplitudes (shaded area denotes SEM), showing two peaks in the *hf_a_*PLV_z_ distribution (35Hz γ and 90Hz *H*γ). **d**, Distribution of average PLV_z_ for all low frequency phases (shaded area denotes SEM), showing two peaks observed in the *lf_p_*PLV_z_ distribution (3Hz, δ and 5Hz, θ).

If CFC facilitates the integration of risk preferences with risk-related reward computations to influence choice behavior, the level of overall OFC CFC may also be related to general risk preference. To test this, we calculated the average δ-θ:γ-*H*γ PLV_z_ for all significant electrodes (*n*=36 electrodes, *n*=10 patients) and tested whether high δ-θ:γ-*H*γ *lf_p_*:*hf_a_* correlated with differences in risk indifference points. We observed that individuals with stronger δ-θ:γ-*H*γ CFC had greater risk-seeking behavior (*R^2^*=0.44, *P*=0.04*, r*=-0.67 Pearson correlation; Extended Data Fig.7b).

### Targeted OFC stimulation causes risk-averse behavior

The representation of inter-individual risk preferences in intra-OFC δ-θ coherence raises the possibility that targeted OFC modulation may be effective in modulating risky decision-making behavior. To test this possibility, and to ascertain the causal role of human OFC in determining risk preferences, we performed intracranial OFC stimulation during gambling task play in a separate, smaller iEEG patient cohort (*n*=3). We delivered continuous, bipolar, high frequency (100Hz, 1-2mA square-waveform), electrical stimulation between the deepest OFC electrode, located in grey matter, and the adjacent, immediately superficial electrode, located in white matter (see Methods). Patients in the stimulation cohort completed two sessions of a shortened version (100 trials) of the gambling task: once in the absence of stimulation (pre-stimulation condition) to characterize baseline risk preferences and a second time during OFC stimulation (stimulation condition). In 2 of these patients, post-hoc anatomical analyses confirmed that stimulation was successfully delivered to a grey matter OFC electrode (Fig.4a,d). In both patients that underwent grey matter stimulation, we observed a robust and consistent modulation of risk-taking behavior, with both patients exhibiting markedly increased risk-averse behavior during stimulation. Specifically, both patients chose to gamble significantly less often (from 55.5 and 47% to 37.0 and 22%, respectively; *P*<0.001, *P*<10E-4 respectively using bootstrapping, see Methods; Fig.4b,e). Consistently, both patients showed lower risk indifference points (from 43.4 and 53.5% to 61.5 and 79.1%, respectively; Fig.4g,h). Because OFC disruption has been associated with impulsivity, we also examined whether stimulation influenced patients’ reaction times. We found that stimulation was associated with faster reaction times in both patients (s01: 0.92±0.3s to 0,85±0.2s, p<0.05, t-test; s02: 2.48±1.46 to 1.5±0.48s, p<10E-8). In contrast, a third patient in which both OFC electrodes were determined post-hoc to be located in white matter we found no modulation or risk preferences or reaction times (Extended Data Fig. 8a).

These results suggest a causal role for OFC in maintaining individual-specific risk preferences and demonstrate that targeted stimulation of the human OFC during risky decision-making results in risk-averse, faster choices.

## Discussion

Individual attitudes in the face of risky decisions vary widely in healthy subjects. These risk preferences are related to a variety of socio-economic factors and have significant importance in behavioral economics (13, 27), but rest upon a strong biological foundation. However, despite neuroanatomical (30) and genetic (31) evidence for the biological basis of risk preferences underlying economic choices, their neurophysiological basis remains poorly understood. This lack of understanding has translational consequences for the numerous psychiatric conditions characterized by altered risky decision-making behavior such as addiction (32), gambling (6), depression (5), and anxiety (33).

Here, we set out to examine the neurophysiological representation of risk preferences in OFC oscillatory activity using intracranial recordings from human patients. We show that risk preferences are associated with subject-specific patterns of low frequency (delta-theta, δ-θ) intra-OFC coherence: patients with higher intra-OFC δ-θ coherence levels engaged in riskier behavior, a relationship that was not observed with δ-θ power modulation (Extended Data Fig.5) and showed anatomical (Extended Data Fig.6) and frequency (Extended Data Fig.4) specificity. In addition, we observed that the phase of δ-θ oscillations modulated the activity of local high frequency (broadband gamma, γ-*H*γ) amplitude (Fig.3) through CFC, whose magnitude also correlated with risk preferences (Extended Data Fig.7b). Finally, direct electrical stimulation of the human OFC modulated risk preferences, causing more risk-averse behavior. Together, we demonstrate a neurophysiological basis for the representation of inter-individual risk preferences in human OFC, propose a model for functional integration of risk-related reward computations through CFC, and show the causal role of intra-OFC neural activity in maintaining functional risk-taking behaviors.

These observations add a new mechanistic layer to the role of OFC in reward-based decision making demonstrated in animal models (8, 9, 11, 21) and humans (34, 35), and expand on recent studies that have started to characterize reward-related neural activity in the human brain using intracranial approaches (12, 26, 36, 37). Specifically, they highlight a central role for δ-θ oscillatory phase in decision-making under uncertainty, adding to previous evidence implicating θ oscillatory activity in decision-making (19, 21, 38, 39). Low frequency oscillations modulate the excitability of neuronal ensembles at coarser temporal and wider spatial scales than high frequency activity, facilitating integration over long temporal scales and large spatial regions (14, 15). We propose that functional communication through δ-θ coherence may act as the organizing principle to facilitate circuit-wide integration of distributed computations into single utility estimates in OFC. During risky decision-making, OFC consolidates multiple decision-relevant computations (risk, win probability, expected value) into a single utility estimate for available decision options (40, 41). This notion is consistent with our observation that patients with higher overall δ-θ coherence engaged in riskier behavior (Fig.2) in our task (Fig.1a). Greater reward-information integration through higher δ-θ coherence may facilitate a more efficient representation of gamble utilities and enhance the salience of risky options. This would, in turn, favor risky choices and result in risk-seeking behavior, diminishing risk-averse biases common in human economic behavior (27). Conversely, safe bets do not require consideration of risk or win probability and are, therefore, less dependent on the functional integration of risk-related reward computations. Consequently, patients with lower intra-OFC δ-θ coherence exhibit risk-aversion, defaulting to the safe bet option that requires minimal utility estimations.

A prediction of this model is that δ-θ phase should interact with the high frequency substrate of reward computations (12). Consistent with this notion, we observed that the phase of δ-θ oscillations modulated the amplitude of γ-*H*γ activity through CFC (Fig.3) in OFC. CFC is a widespread neural phenomenon (16, 42) through which circuit-level networks operating at slow (behavioral) timescales interact with the fast cortical processing widely present in the human prefrontal cortex and other regions (16, 28). Despite its involvement in multiple cognitive processes including memory (43), spatial navigation (44) and decision-making behavior (21, 29), the computational role of CFC is not fully established. Our results suggest that, during risky decision-making, OFC CFC may integrate relatively stable risk preferences, reflected in circuit-wide oscillatory coherence, with fast, decision-relevant computations, reflected in local high frequency amplitude (Fig.5). This notion is further supported by our observed relationship between δ-θ:γ-*H*γ CFC and risk preferences (Extended Data Fig.7a). Our observations are limited to a specific behavioral context and a single relevant brain region, but they open the door to examining whether similar interactions between oscillatory coherence, CFC and local high-frequency activity may be a generalizable mechanism to achieve integration of stable, trait-like preferences with transient activations related to ongoing task and behavioral context. If so, neurophysiological coherence estimates may provide a normative metric for the neural basis of natural variation in human behavior.

Finally, our results provide causal evidence for the role of the human OFC in decision-making and demonstrate the potential for precise behavioral modulation through neurostimulation. Targeted OFC stimulation shifted individual risk preferences, causing patients to show more risk-averse behavior compared to their pre-stimulation baseline (Fig.4). This effect was limited to the modulation of OFC grey matter activity (Fig.4), as opposed to stimulation of white matter which had no effect on risk-taking behavior (Extended Data Fig.8a). Our observations are consistent with previous work reporting a lack of behavioral changes with white matter stimulation (45). This finding suggests that these effects are specific to direct modulation of OFC activity, rather than adjacent fibers of passage (46). Finally, OFC stimulation also resulted in faster reaction times across both patients (Fig.4). Prior studies have shown that OFC lesions increase impulsive behavior, characterized by fast reaction times (47). Therefore, our results are consistent with the notion that high-frequency stimulation results in a temporary inactivation of OFC akin to a reversible lesion, and with prior proposals that impulsivity (reaction times) and risk preferences may be separable behaviorally and neurally.

**Figure 4.**
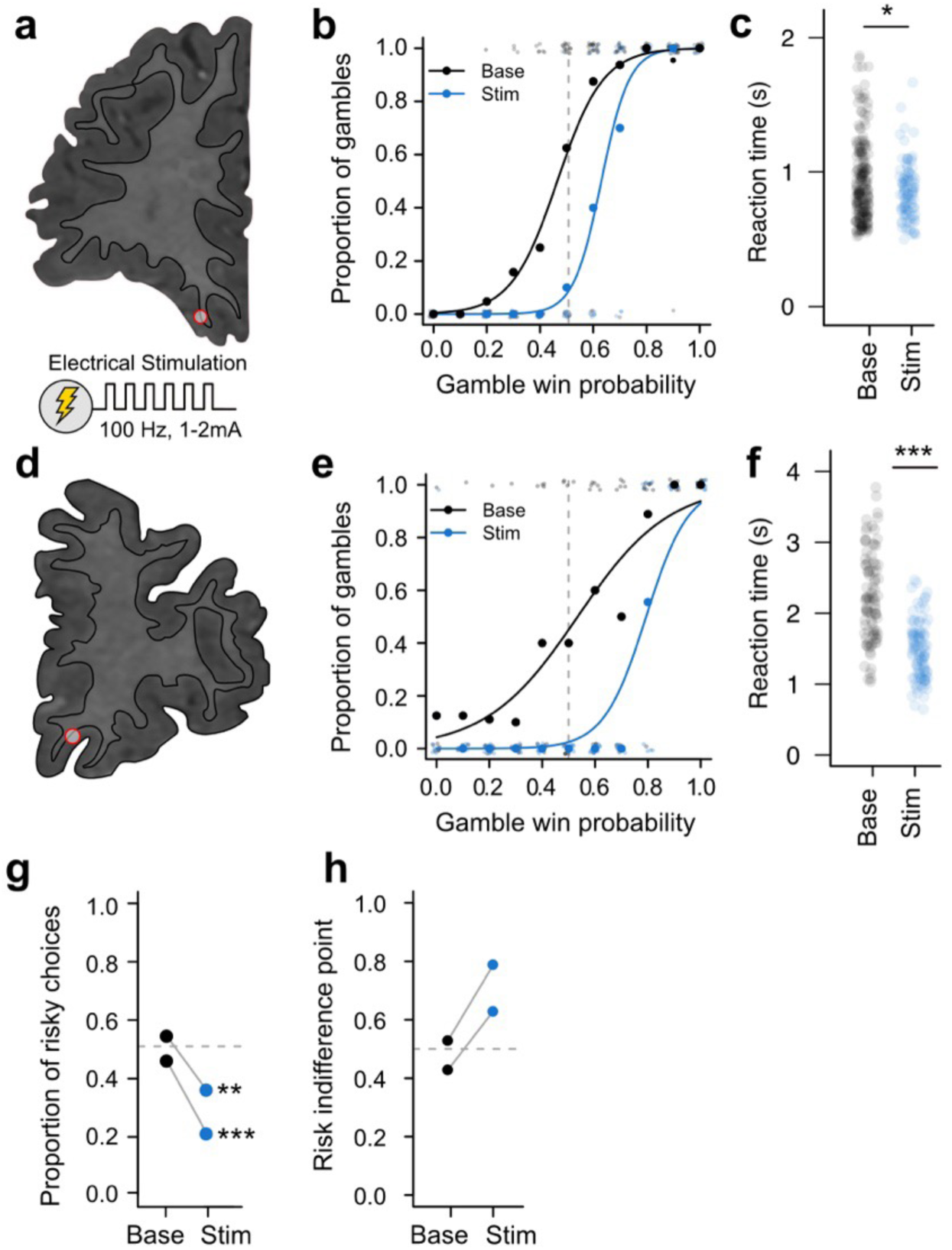
Orbitofrontal cortex stimulation modulates risk preferences and reaction times. **a**, Anatomical location of electrodes used for stimulation in patient s01. The two deepest electrodes were selected for stimulation, and their anatomical location was subsequently verified by examination of the co-registered MR-CT images. The deepest electrode, located in grey matter, was used as the anode (red point), and the second deepest electrode, located in white matter, as the cathode. **b**, Behavioral risk preference modulation during stimulation in patient s01. Plot shows the proportion of gambles that the patient chose as a function of the win probability of the proposed gamble, in 10% win probability increments. The patient completed the behavioral task twice: once without stimulation (Pre-stim condition, black line) and a second time during continuous OFC stimulation (Stim condition, blue line). The leftward shift of the fitted curve indicates a change towards risk-averse behavior. Smaller grey points indicate trial-by-trial choices (0=safe bet choice; 1=gamble choice). **c,** Reaction times in patient s01. Reaction times (s) in individual trials were slower during baseline (Base, grey points) than during stimulation (Stim, blue points) conditions. **d-f**, As **a-c**, but for patient s02. **g**, Proportion of risky choices for both patients across conditions: Pre-stim (black points) and Stim (blue points). The proportion of risky choices was significantly decreased in the stimulation condition for both patients (both *P*<0.01, bootstrapped). **h**, As **g**, but for risk indifference point.

**Figure 5.**
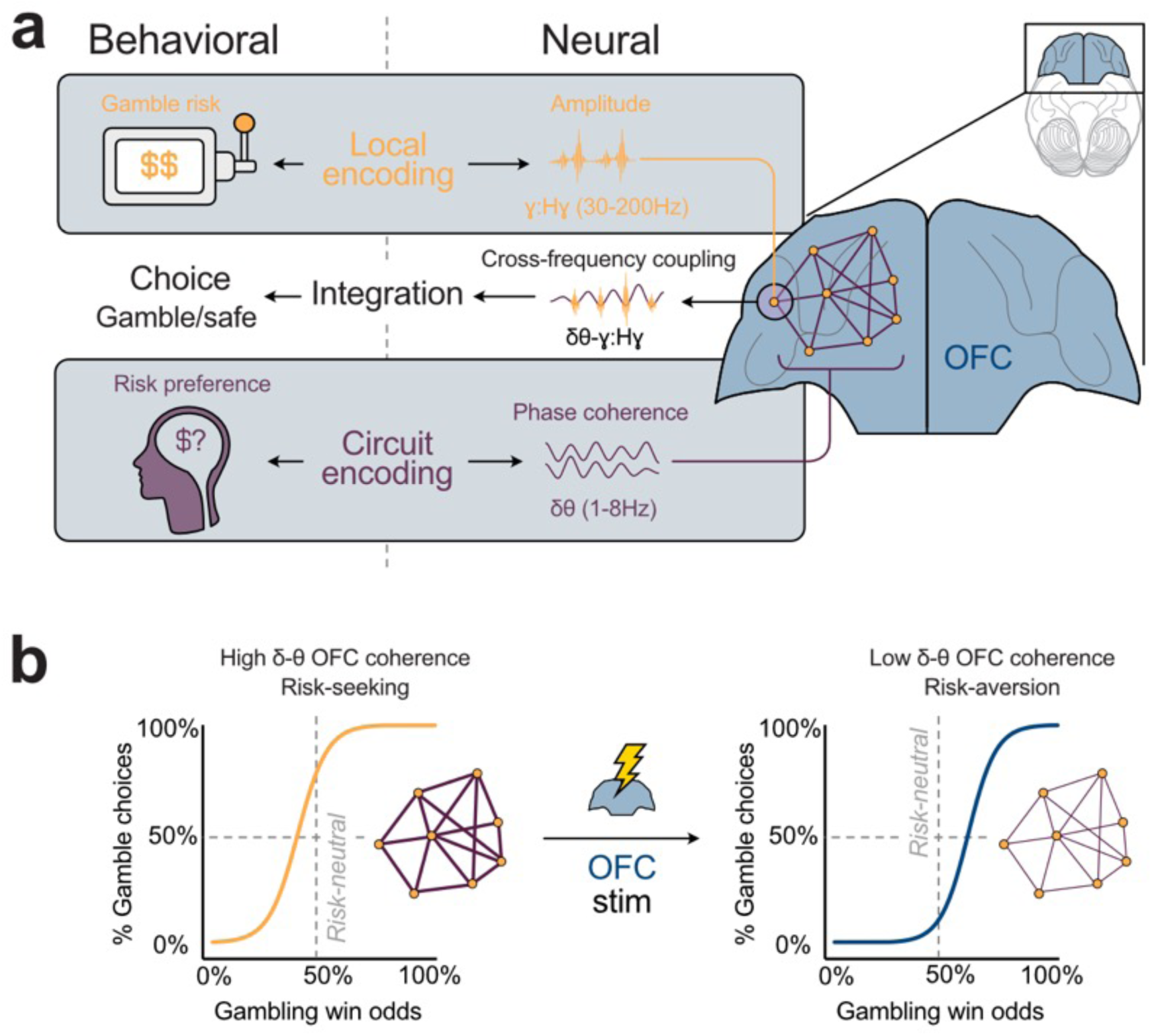
The neurophysiological basis of risk preferences and behavior in human OFC. **a**, Conceptual model. Behaviorally (left), both the characteristics of the external gamble (i.e. risk) and the player’s internal risk preferences need to be integrated to generate adaptive choices. Reward related computations, including gamble risk, are represented in local broadband gamma (γ*-H*γ, 30-200Hz) activity across distributed OFC sites (local encoding, yellow, top). Conversely, inter-individual risk preferences are reflected by circuit-wide functional coordination within OFC, reflected in δ-θ (1-8Hz) coherence across OFC sites (circuit encoding, purple, bottom). Phase-amplitude cross frequency coupling between δ-θ phase and γ*-H*γ amplitude facilitates functional integration of risk preference and reward computations, both neurally and behaviorally (middle). **b**, Targeted OFC stimulation modulates inter-individual risk preferences. Patients with high OFC δ-θ coherence are more risk-seeking and choose risky gambles more often (left), whereas patients with low δ-θ coherence are more risk-averse. Targeted OFC stimulation disrupts δ-θ coherence, local γ*-H*γ encoding and/or CFC-mediated integration, favoring safe choices that are less dependent on functional integration of multiple reward computations and inducing risk-averse behavior.

Prior evidence has shown that OFC microstimulation is effective at modulating choice behavior during reward-based decision-making in non-human primates (48). Our results complement these observations by showing the effect of continuous macrostimulation during risky decision-making in the human OFC. Here, stimulation is likely to disrupt the OFC dynamics necessary for risky choice by altering the encoding of reward computations (risk, win probability) in high frequency power (12, 26), interfering with stable preference representations in intra-OFC coherence, or impeding cross-frequency integration through CFC (28). Consequently, patients would default to safe bet choices that do not require the successful integration of multiple reward-related computations, as we observed (Fig.4).

The behavioral changes caused by targeted OFC stimulation were behaviorally precise. Targeted OFC stimulation specifically modulated risk preferences, but did not abolish reward or win probability sensitivity, suggesting that targeted modulation of individual behavioral preferences without severe disruption of behavior is possible. Since the patients in our sample exhibited risk preferences that were approximately optimal for our task (Extended Data Fig.1d), shifting their behavior to be more risk-averse resulted in overall less optimal choices. However, this stimulation strategy may be capable of correcting behavioral deficits present in psychiatric conditions where decision-making behavior, and likely its underlying neurobiological substrates, are altered. More generally, our results provide a potential transdiagnostic biomarker for altered risk preferences in psychiatric pathologies, which our results predict will be accompanied by changes in the degree of intra-OFC δ-θ coherence, an idea that could be tested by studying intracranial patients with relevant co-morbidities (49).

In addition, OFC stimulation emerges as a candidate neurostimulation treatment for individuals with pathological risk-taking behaviors, particularly those involving heightened risk-seeking (2, 5, 6). Our results also highlight the utility of neuromodulatory strategies to treat specific pathological behavior, rather than using task-free stimulation strategies (50), and the value of symptom-specific, transdiagnostic therapeutic approaches focused on behavioral modulation (7). Conceptually, the development of closed-loop neurostimulation approaches that are sensitive to behavioral context and neural states may be especially relevant (7, 39, 48). Combined with recent evidence that OFC stimulation modulates affective states (51), there is growing support for stimulation-based therapies specifically targeting the OFC. Lastly, whereas our high frequency stimulation strategy resulted in an increase in risk-aversion (Fig.4), our model (Fig.5) supports the notion that a different stimulation strategy designed to entrain OFC activity to a δ-θ rhythm(52), shown to affect decision-making (39), could achieve opposite effects (i.e. an increase in risk-seeking), providing an alternative therapeutic route for disorders characterized by risk-aversion such as depression (5).

In summary, we defined a novel neurophysiological substrate of inter-individual risk preferences in the human OFC and provide causal evidence for behavioral modulation through targeted neurostimulation. Mechanistically, we proposed that intra-OFC oscillatory coherence, reflecting stable risk preferences, facilitates functional integration of multiple reward computations into single utility estimates through CFC. Modulation of these processes through direct OFC stimulation resulted in robust behavioral changes, markedly decreasing risk-seeking behaviors, while maintaining overall reward sensitivity. Our findings demonstrate the utility of targeted OFC stimulation for modulating risky behaviors and open the door to developing of novel, transdiagnostic neurostimulation interventions to correct pathological risk preferences present in psychiatric conditions such as addiction and gambling disorders.

## Materials and methods

### Patients

iEEG data was collected from 15 (5 female) adult subjects with intractable epilepsy who were implanted with chronic subdural grid and/or strip electrodes as part of a pre-operative procedure to localize the epileptogenic focus. Stimulation datasets were collected in an additional *n*=3 adult epilepsy patients. We paid careful attention to the patient’s neurological condition and only tested when the patient was fully alert and cooperative. The surgeons determined electrode placement and treatment based solely on the clinical needs of each patient. Patient recordings took place at University of California and at Mount Sinai West. As part of the clinical observation procedure, patients were off anti-epileptic medication during data collection. The time frame available for data collection (days) limited our ability to assess the stability of risk preferences across longer timescales. Healthy participants (*n*=20) with no prior history of neurological disease played an identical version of the gambling task. All subjects gave written informed consent to participate in the study in accordance with the University of California and the Mount Sinai Institutional Review Boards.

### Gambling task

Patients played a gambling game in which they chose between a $10 safe bet and a higher payoff gamble (e.g. $30), as previously described(12). For our behavioral control analyses, we collected behavioral data from *n*=20 healthy volunteers. For full details on task design, see Supporting Information.

### Stimulation bootstrapping

To estimate the significance of behavioral modulation during stimulation, we compared the proportion of risky choices in baseline versus stimulation epochs using a bootstrapping approach.

First, we sampled the proportion of risky choices under the baseline condition by selecting *n*=50 patient choices (without replacement) during the baseline condition and calculating the resulting risky choice percentage. We repeated this procedure 10,000 times to generate a null distribution. Finally, we compared the Null distribution to the observed proportion of risky choices under the stimulation condition to obtain a permutation p-value, equal to the proportion of shuffles showing a more extreme value than the experimentally observed one.

### Prospect theory model

We used a prospect-theory derived computational model to estimate model-based risk preferences in our patient sample, and to simulate choice behavior from artificial agents with different risk preferences and determine optimal risk attitude (i.e. the risk preference that maximizes profit) in our gambling task. For full details, see Supporting Information.

### iEEG electrode localization

Full details on electrode localization methods are available in earlier publications (12, 53). Location of hippocampal regions (Fig. S7) was determined similarly, limiting analyses to electrodes in hippocampal grey matter. Electrodes within 3cm of clinically identified seizure focus or abnormal tissue (e.g. cortical dysplasia) are excluded from analyses. Our neurosurgical iEEG approach limited our investigation to clinically targeted regions, impeding the investigation of other regions likely to be implicated in risk encoding, such as the insula and striatum (54).

### iEEG recording and preprocessing

iEEG electrode number and location depends strictly on clinical strategy. iEEG LFP activity was recorded and stored with behavioral data: 64-256 channels amplified x10000, analog filtered (0.01-1000 Hz) with a minimum 1KHz digitization rate, re-referenced to a single white matter electrode off-line, and high-pass filtered above 1Hz with a symmetrical (phase true) finite impulse response (FIR) filter (∼35 dB/octave roll-off). Epileptic electrodes and epochs were discarded after careful visual examination of the collected datasets (CT/MRI and iEEG) by a neurologist or a researcher. We used manual and automated pre-processing methods to exclude movement-related and epileptiform activity. Channels with low signal-to-noise ratio were identified and removed (i.e. 60 Hz line interference, electromagnetic noise from hospital equipment, amplifier saturation, poor contact with cortical surface. All electrodes and trials were visually examined and trials in which movement artifacts or epileptiform activity were identified were excluded from analyses. LFP data was preprocessed: low-pass filtered at 200Hz, high-pass filtered at 0.1Hz, notch filtered at 60Hz and harmonics and down-sampled to 1KHz if necessary. LFP recordings were trialed around events of interest (e.g. baseline or decision epochs, Fig. 1B).

### Coherence and linear regression analyses

The raw LFP signal for every OFC electrode across patients was decomposed into time-frequency representations using the fast Fourier transform with a single Hanning taper (using Fieldtrip toolbox ft_freqanalysis function). Two epochs were used to compute coherence: the baseline epoch (500 ms following trial reveal) and the decision epoch (1000ms preceding button press for choice). The oscillatory synchrony between pairs of electrodes was quantified as the coherence coefficient (using Fieldtrip toolbox ft_connectivityanalysis function) (55). Coherence was calculated between all pairs of OFC and hippocampus electrodes (Fig. S7) within subject for delta (δ, 1-4Hz), theta (θ, 5-8Hz), delta-theta (δ-θ, 1-8Hz), alpha (α, 9-12Hz), and beta (β, 13-30Hz) bands with a 1Hz frequency resolution. To evaluate the association between risk preference and coherence, we conducted a linear regression analysis to assess the relationship between an individual subject’s risk indifference point and their mean coherence value between all electrode pairs for each frequency band.

### Cross Frequency Coupling analyses

We used cross frequency analyses to estimate the impact of low-frequency neural oscillatory phase on high-amplitude power. Full details about CFC methods, as well as phase-locking value statistical methods, are available in the Supplementary Materials.

### Electrical stimulation

Patients in the stimulation cohort played the gambling task twice: first, in the absence of stimulation, to evaluate baseline risk preference, and a second time under continuous electrical stimulation of OFC sites as above. After a first stimulation-free behavioral dataset has been collected, the patient completed a second task block during stimulation to assess the impacts of stimulation on behavior. To minimize task duration and allow for collection of both baseline and stimulation datasets, we halved the number of trials (*n*=100), with the trial subset spanning the same range of win probability and risk as the full behavioral task. Brain stimulation was performed after enough seizure data had been collected and anticonvulsant medications have been restarted, to minimize the possibility of triggering a seizure. None of the electrodes stimulated were located within 3cm of known or suspected epileptogenic zones(^56^). Stimulation was carried out using a Natus Cortical Stimulator that allows bipolar single-electrode stimulation. We applied bipolar stimulation and delivered individual charge-balanced pulses on single grey matter electrodes (as confirmed post-hoc by anatomical localization and the closest white matter electrode in the same iEEG surgical tract). We used 1 or 2mA amplitude for stimulation, well below the 6mA routinely used during cortical mapping in epilepsy, to minimize the likelihood of triggering seizures. To minimize the chance of triggering epileptic seizures, we first verified that none of the electrodes selected for stimulation were located within 3cm of known or suspected epileptogenic zones and second carried out initial single-pulse stimulation and monitored for epileptic after-discharges; no after-discharges were visible in any electrodes. We applied stimulation using biphasic, constant-current trains of stimulation pulses at 100 Hz, with 100µs pulse width. We assessed the impact of stimulation by examining patients’ behavior during stimulation and comparing risk attitudes (risk indifference points) it to pre-stimulation baseline.

## Acknowledgements and funding sources

We would like to thank L. Nuñez and D. Bolden for help with data collection, patients for their willingness to participate in this research, the clinical team for support with data collection and Peter Rudebeck, Erin Rich, Craig Mermel and Salman Qasim for fruitful discussions. The project described was supported by the National Institutes of Mental Health (grants number K01MH108815 and R01MH124763 to I.S.) and the National Science Foundation (graduate research fellowship to A.S.).

## Notes

**Competing Interest Statement:** the authors declare no competing interests.

### Competing Interest Statement

The authors have declared no competing interest.

### Summary of Updates

Incorrect author name generated by BiorXiv autofill feature. Author Lu Jin has been corrected from Ju Lin.

