## Supplemental Materials for "Neuromodulation of risk preferences encoded in human orbitofrontal cortex activity"

**This PDF file includes:**

Supporting text

Figures S1 to S8

Table S1

Supporting Information Text

**Extended methods.**

**Materials and methods**

### Gambling task

Patients played a gambling game in which they chose between a $10 safe bet and a higher payoff gamble (e.g. $30), as previously described (12). Gamble win probability varied parametrically round by round: subjects are shown a number between 0-10, at which point they choose whether to keep the safe bet prize or the risky gamble. Patients had up to 4s to make a choice, or a timeout occurred. 550ms after a selection is made, a second number (also 0-10) is revealed. Both numbers are drawn from a uniform probability distribution, and patients have full knowledge of the game. If the second number is greater than the first one, the gamble results in a win. Risk is maximal at 50% win probability (shown number = 5) and lowest at win probability = 0% (shown number = 10) and win probability = 100% (shown number = 0). The game was designed to allow for variation in risk preferences. To assess risk preferences, we fit a sigmoidal curve over patient’s choices binned over 10% win probability bins and identify the risk indifference point at which the patient is equally likely to choose a gamble or a safe bet. Patients played a total of 200 rounds (plus practice rounds), and a full experimental run typically lasted 12-15min. Location of safe bet and gamble options (left/right) was randomized across trials. Patients completed a training session prior to the game in which they played at least 10 rounds under the experimenter’s supervision until they understood the task, at which point the game was started. This gambling task minimized other cognitive demands (working memory, learning, etc.), while including variation in variables related to decision-making including win probability and risk.

*Stimulation bootstrapping*

To estimate the significance of behavioral modulation during stimulation, we compared the proportion of risky choices in baseline versus stimulation epochs using a bootstrapping approach. First, we sampled the proportion of risky choices under the baseline condition by selecting *n*=50 patient choices (without replacement) during the baseline condition and calculating the resulting risky choice percentage. We repeated this procedure 10,000 times to generate a null distribution. Finally, we compared the Null distribution to the observed proportion of risky choices under the stimulation condition to obtain a permutation p-value, equal to the proportion of shuffles showing a more extreme value than the experimentally observed one.

**Prospect theory model**

We used a prospect-theory derived computational model to estimate model-based risk preferences in our patient sample, and to simulate choice behavior from artificial agents with different risk preferences and determine optimal risk attitude (i.e. the risk preference that maximizes profit) in our gambling task. For every simulated risk attitude, we simulated 500 sessions of 200 trials using the same gambling task structure played by iEEG patients. In the prospect theory framework, an individual’s decisions are influenced by the value of their choices as well as their subjective aversion to risky decisions. Therefore, prospect theory models of risky decision behavior transform choice values to subjective utilities using a constant risk parameter, which represents the curvature of the individual’s utility function(29). For every trial, safe bet and gamble payoff values were converted to subjective utilities using the simulated agent’s risk parameter value and the gamble win probability:

$$U(gamble) = P(win)\times{V(\mathrm{gamble})}^{risk}$$

$$U(safe) = {V(safe)}^{risk}$$

Where $P(win)$ equals the win probability, $V(\mathrm{gamble})$ equals the potential payoff value of the gamble, and $V(safe)$ equals the value of the safe bet (always $10). The values of the gamble and safe offers are weighted by the exponent, *risk*, which equals the known risk attitude of the artificial agent. The simulated risk parameter values ranged from 0.1 to 2.5 to reflect risk-averse (*risk* < 1), risk-neutral (*risk* = 1), and risk-seeking (*risk* > 1) behaviors(2). The resulting utilities, $U(gamble)$ and $U(safe)$, were then used to estimate the agent’s choice probability with a softmax rule with a fixed exploration parameter, $\beta$, equal to 5 for all simulations:

$$P(gamble) = \frac{exp(\beta\times u(gamble))}{exp(\beta\times u(gamble)) + exp(\beta\times u(safe))}$$

The risk indifference point was estimated for each risk parameter using the average simulated choice behavior to fit a sigmoidal curve over the agent’s choices to find the value of $P(win)$ where $P(gamble)$= 0.5. To validate that the prospect theory risk parameter reflected the same risk attitude as the risk indifference point, which was used to quantify risk attitudes for iEEG patients, the value of the risk parameter and risk indifference point for each simulation were correlated (Fig S1A, Pearson r = -0.98, *p* = 1.9e-33). To determine the optimal risk attitude, final profits for all simulations for each risk parameter value were averaged to give one estimate of the average total profit for every risk parameter (Fig S1B, C).

**Cross Frequency Coupling analyses**

*Computing Cross Frequency Coupling Using Phase Locking Values*

Cross frequency phase-amplitude coupling was calculated as the phase locking value (PLV) between two signals of interest: a low frequency phase (*lf_p_*) and a high frequency amplitude (*hf_a_*). The low frequencies for phase (*lf_p_*, 2-20Hz in 1 Hz steps) were delta-theta (δ-θ, 2-8Hz), alpha (α, 8-12Hz), and slow-beta (β, 12-20Hz) low frequency bands. The high frequencies for amplitude (*f_a_*, 5-200Hz in 5Hz steps) were theta (θ, 5Hz), alpha (α, 10Hz), beta (β, 15,20,25Hz), low-gamma (γ, 30-60Hz), and high-gamma (*H*γ, 60-200Hz). Low and high-gamma were combined into broadband gamma for analyses (γ-*H*γ, 30-200Hz). We used an exploratory approach(56) to calculate the PLV between every phase (*lf_p_*) and amplitude (*hf_a_*) frequency combination, resulting in 760 phase locking values spanning multiple cross-frequency interactions within each OFC electrode.

The PLV was calculated between the low frequency phase time series, $\phi_{{lf}_{p}}$, and the low frequency phase of the high frequency amplitude, $\phi_{{hf}_{a}}$, for each *lf_p_*:*hf_a_* pair. The low frequency phase, $\phi_{{lf}_{p}}$ and high frequency amplitude envelope were computed using gaussian chirplet transforms^16^, and the angle of the time domain of the high frequency amplitude envelope was used to extract the phase time series, $\phi_{{hf}_{a}}$. Phase-amplitude coupling was estimated only for decision epochs. The decision epochs (-1 to 0 seconds prior to choice) were extracted for every trial from $\phi_{{lf}_{p}}$ and $\phi_{{hf}_{a}}$respectively and combined to estimate a single coupling value across trials. Then, the PLV was computed between $\phi_{{lf}_{p}}$and $\phi_{{hf}_{a}}$, giving an estimate of the strength of synchronization between the phase of the low frequency *lf_p_* and amplitude of the high frequency *hf_a_* during the decision epoch^93–97^. The *lf_p_*:*hf_a_* PLV is defined as:

$${PLV}_{{lf}_{p}:{hf}_{a}}= \left| \left\langle exp(i[\phi_{{lf}_{p}}-\phi_{{hf}_{a}}]) \right\rangle\right|$$

Where $\left\langle\cdot\right\rangle$ denotes mean of the *lf_p_*:*hf_a_* phase difference estimates across timepoints and $\left| \cdot\right|$ denotes the magnitude of the PLV*_lfp_*_:_*_hfa_* vector, or the strength of the phase locking between $\phi_{{lf}_{p}}$and $\phi_{{hf}_{a}}$. PLV can range from 0 to 1, where PLV = 0 indicates two frequencies are completely desynchronized and PLV = 1 indicates perfect synchronization.

*Individual PLV Normalization and Significance Testing*

For each PLV*_lfp_*_:_*_hfa_*, the phase locking statistic must be computed to assess whether the observed PLV was significantly greater than phase locking due to random chance. The standard practice for estimating phase locking statistics is to use nonparametric statistical testing using a randomized permutation distribution as the null hypothesis for each PLV*_lfp_*_:_*_hfa(57)_*. We computed a null distribution of 1000 surrogate (N*_surr_*) values for each *lf_p_*:*hf_a_* combination. To calculate each PLV*_surr_*, $\phi_{{lf}_{p}}$was circularly shifted by a random number of samples and the PLV between the shifted $\phi_{{lf}_{p}}$and original $\phi_{{hf}_{a}}$was recalculated. The random shift was resampled for every surrogate iteration to generate 1000 PLV*_surr_* values. If the observed PLV*_lfp_*_:_*_hfa_* was not just the result of random chance, it should be consistently greater than the PLV*_surr_* estimates. The permutation *p*-value was then estimated as of the proportion of PLV*_surr_* larger than PLV*_lfp_*_:_*_hfa_*:

$$p= \frac{{\#}_{i=1}^{N_{surr}}[{PLV}_{surr}>{PLV}_{{lf}_{p}:{hf}_{a}}]}{N_{surr}}$$

In addition to providing a significance estimate, the PLV*_surr_* distribution was used to normalize the observed PLV*_lfp_*_:_*_hfa_* to allow for comparison of PLVs across *lf_p_*:*hf_a_* pairs within and between electrodes (*n*=130). The normalized PLV*_lfp_*_:_*_hfa_* was transformed into a *z*-score using the mean and standard deviation of its PLV*_surr_* distribution.

$${PLV}_{z}= \frac{{PLV}_{{lf}_{p}:{hf}_{a}}- [\mu({PLV}_{surr})]}{[\sigma({PLV}_{surr})]}$$

This process was repeated for all combinations of *lf_p_*:*hf_a_* (*n*=760 pairs) for each OFC electrode (*n*=130). The multiple cross-frequency interactions were visualized as a comodulogram(16), where the magnitude of the ${PLV}_{z}$ for all *lf_p_*:*hf_a_* pairs was represented on the z-axis (Fig 3A, single electrode example).

*Electrode-Level PLV Significance Testing*

To determine whether significant cross-frequency phase-amplitude coupling was present at the electrode-level, we sought to determine whether significant clusters of adjacent *lf_p_*:*hf_a_* pairs exceeded the surrogate significance thresholds. First, each individual *lf_p_*:*hf_a_* *p*-value was corrected to account for multiple comparisons. There were 760 highly dependent significance tests performed for every electrode, making the Bonferroni correction approach too stringent. The Benjamini & Hochberg False Discovery Rate (FDR) correction procedure was best suited to reduce the proportion of Type I errors of these dependent data without reducing power of the statistical test(58). PLV *p*-values were corrected on an electrode-level using the FDR-controlling procedure (α=0.05) for dependent tests. Individual *lf_p_*:*hf_a_* pairs were considered significantly coupled if their *FDR Corrected* *p*-value was less than 0.05.

The exploratory nature of our approach requires further statistical thresholding to interpret cross-frequency phase-amplitude synchronization between electrodes. For a given electrode, only a portion of frequency pairs will meet significance criteria, making it difficult to determine a group-level threshold for whether an electrode exhibits significant phase-amplitude coupling. Electrodes were considered significant only if they had at least one significant cluster of *lf_p_*:*hf_a_* pairs. Clusters were estimated using the MATLAB contourf function to group PLV*_z_* pairs in the electrode comodulogram. Cluster-level statistics were computed as the mean PLV*_z_* (Z-statistic). For a cluster to have been extracted from the comodulogram, its cluster statistic was greater than two, corresponding to >2 standard deviations above the chance-level surrogate distributions. Additionally, a cluster contained at least two adjacent *hf_a_* frequencies to exceed the minimum cluster-size threshold. Our cluster-level statistical significance criteria required that at least 95% of a cluster’s corresponding FDR-Corrected *p*-values be below 0.05. For an electrode to have significant cross-frequency phase-amplitude coupling, the electrode had at least one cluster that met all cluster-level criteria (*size:* contains>=2 *hf_a_*, *magnitude:* mean PLV*_z_*>2, *significance threshold:* at least 95% FDR-Corrected *P*<0.05). This cluster-based statistic approach was well-suited for interpreting meaningful multi-frequency phase-amplitude coupling relevant to individual-level behavioral characteristics.

### Electrical stimulation

Patients in the stimulation cohort played the gambling task twice: first, in the absence of stimulation, to evaluate baseline risk preference, and a second time under continuous electrical stimulation of OFC sites as above. After a first stimulation-free behavioral dataset has been collected, the patient completed a second task block during stimulation to assess the impacts of stimulation on behavior. To minimize task duration and allow for collection of both baseline and stimulation datasets, we halved the number of trials (*n*=100), with the trial subset spanning the same range of win probability and risk as the full behavioral task. Brain stimulation was performed after enough seizure data had been collected and anticonvulsant medications have been restarted, to minimize the possibility of triggering a seizure. None of the electrodes stimulated were located within 3cm of known or suspected epileptogenic zones^(59)^. Stimulation was carried out using a Natus Cortical Stimulator that allows bipolar single-electrode stimulation. We applied bipolar stimulation and delivered individual charge-balanced pulses on single grey matter electrodes (as confirmed post-hoc by anatomical localization and the closest white matter electrode in the same iEEG surgical tract). We used 1 or 2mA amplitude for stimulation, well below the 6mA routinely used during cortical mapping in epilepsy, to minimize the likelihood of triggering seizures. To minimize the chance of triggering epileptic seizures, we first verified that none of the electrodes selected for stimulation were located within 3cm of known or suspected epileptogenic zones and second carried out initial single-pulse stimulation and monitored for epileptic after-discharges; no after-discharges were visible in any electrodes. We applied stimulation using biphasic, constant-current trains of stimulation pulses at 100 Hz, with 100µs pulse width. We assessed the impact of stimulation by examining patients’ behavior during stimulation and comparing risk attitudes (risk indifference points) it to pre-stimulation baseline.

### iEEG electrode localization

Full details on electrode localization methods are available in earlier publications^83,87^. Electrode placement during surgery was determined solely on clinical grounds and confirmed post-surgery after visual examination of each patients’ anatomical scans. Briefly, post-operative CT images showing location of individual electrodes were co-registered with preoperative structural MRIs using normalized mutual information algorithms implemented in Fieldtrip^87^. This allowed us to determine the location of each electrode in MR-determined anatomy. Electrode anatomical location is assigned visually by contrasting electrode localizations in the co-registered CT-MR image with surgical notes and electrophysiology files. We focused our main analyses on electrodes determined to be in grey matter in the orbitofrontal region, as identified verified by automatic querying of anatomical Atlases (AAL, AFNI, JuBrain, and Yale Brain Atlas^88^) and confirmed by visual examination by a neurologist or researcher.

Fig. S1.

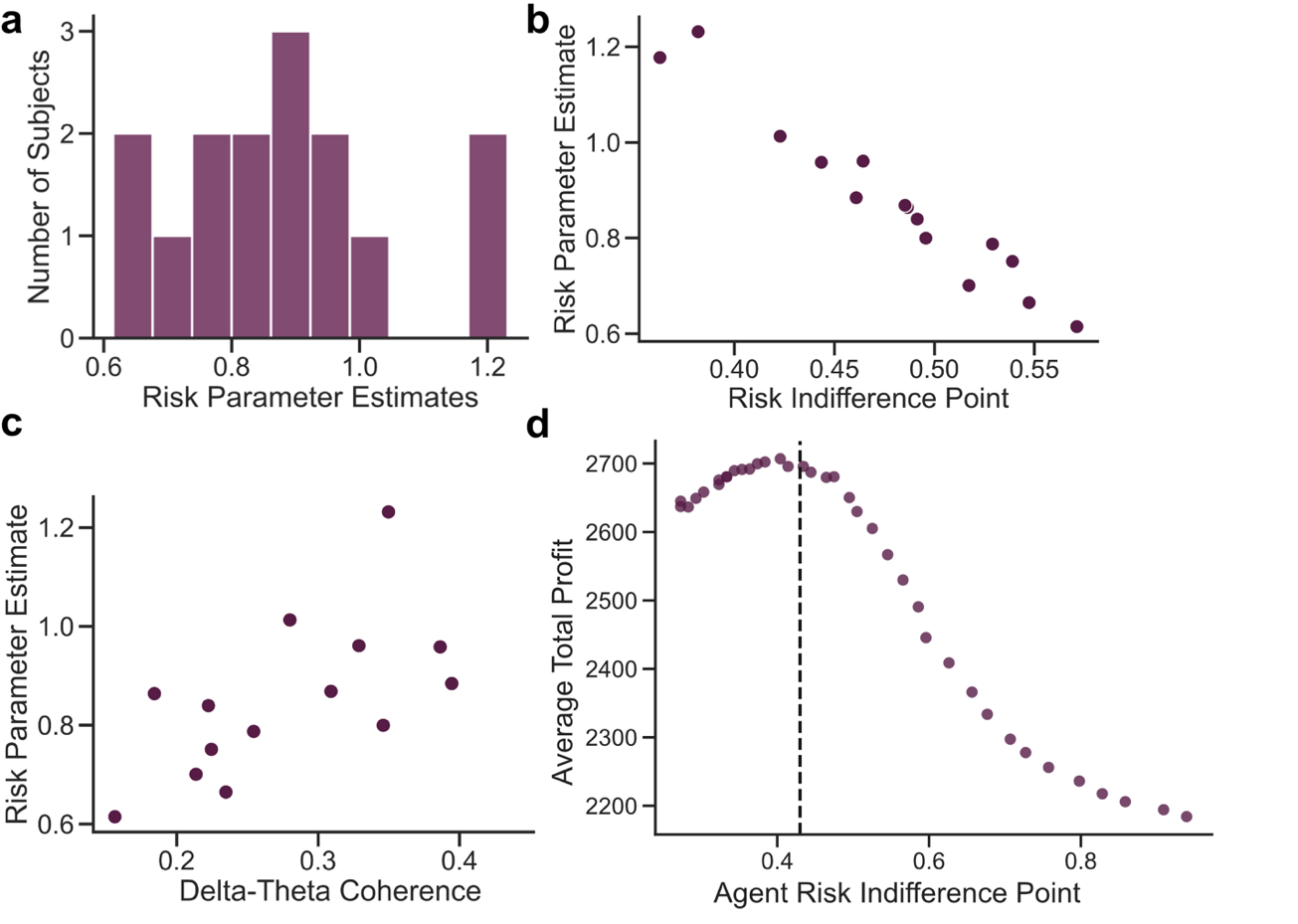
Extended Data Figure 1. Computational Modeling of Risk Preferences. We used a prospect theory model approach to determine optimal risk behavior in the gambling task. a, Distribution of risk parameter values for iEEG subjects (*n*=15). b, Correlation of patients’ model-free risk indifference points and model-based risk parameter estimate (Pearson *r*=0.97, *P*=1.1e-09). c, The prospect theory-derived risk aversion parameter correlates with intra-OFC delta-theta coherence (*R^2^*=0.49, *P*=0.002). d, Optimal choice behavior for our gambling task should maximize profits. The simulated risk parameter values were plotted against their average total profit to determine which risk attitude resulted in the greatest profit. The optimal agent had an approximately risk neutral indifference point of 0.43.

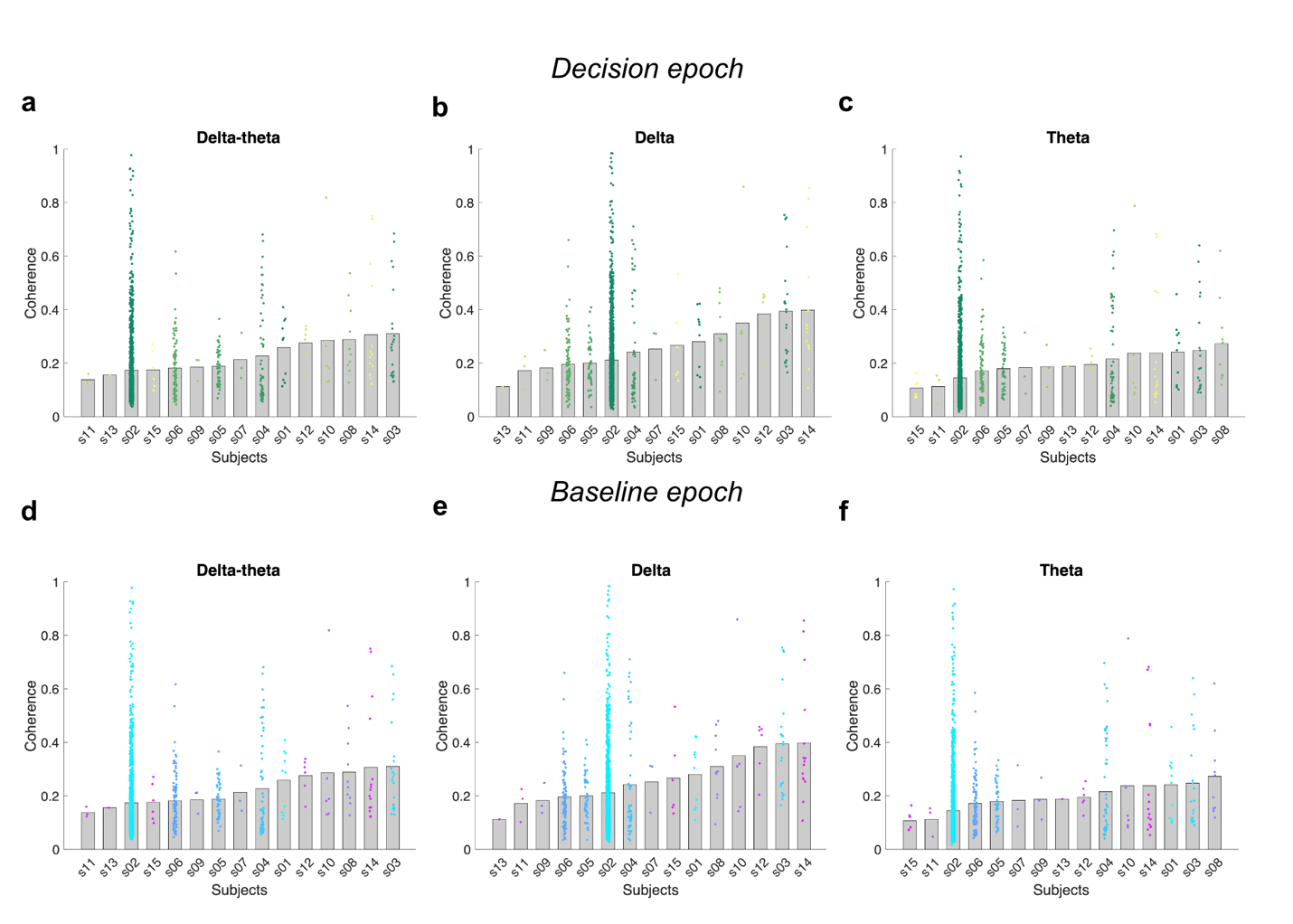
Fig. S2

**Extended Data Figure 2. Coherence of all OFC electrode pairs in delta-theta, delta, and theta bands, grouped by subjects in the decision epoch (a,c) and baseline epoch (d-f).** Dots represent pairwise coherence between pairs of OFC electrodes within each subject; bars represent the mean coherence of each subject. The mean delta-theta coherence for all subjects in decision epoch is 0.29±0.087 and baseline epoch is 0.22±0.058. Mean delta coherence for all subjects in decision epoch is 0.32±0.107, and baseline epoch is 0.27±0.084. Mean theta coherence for all subjects in decision epoch is 0.27±0.084 and baseline epoch is 0.19±0.050.

Fig. S3.

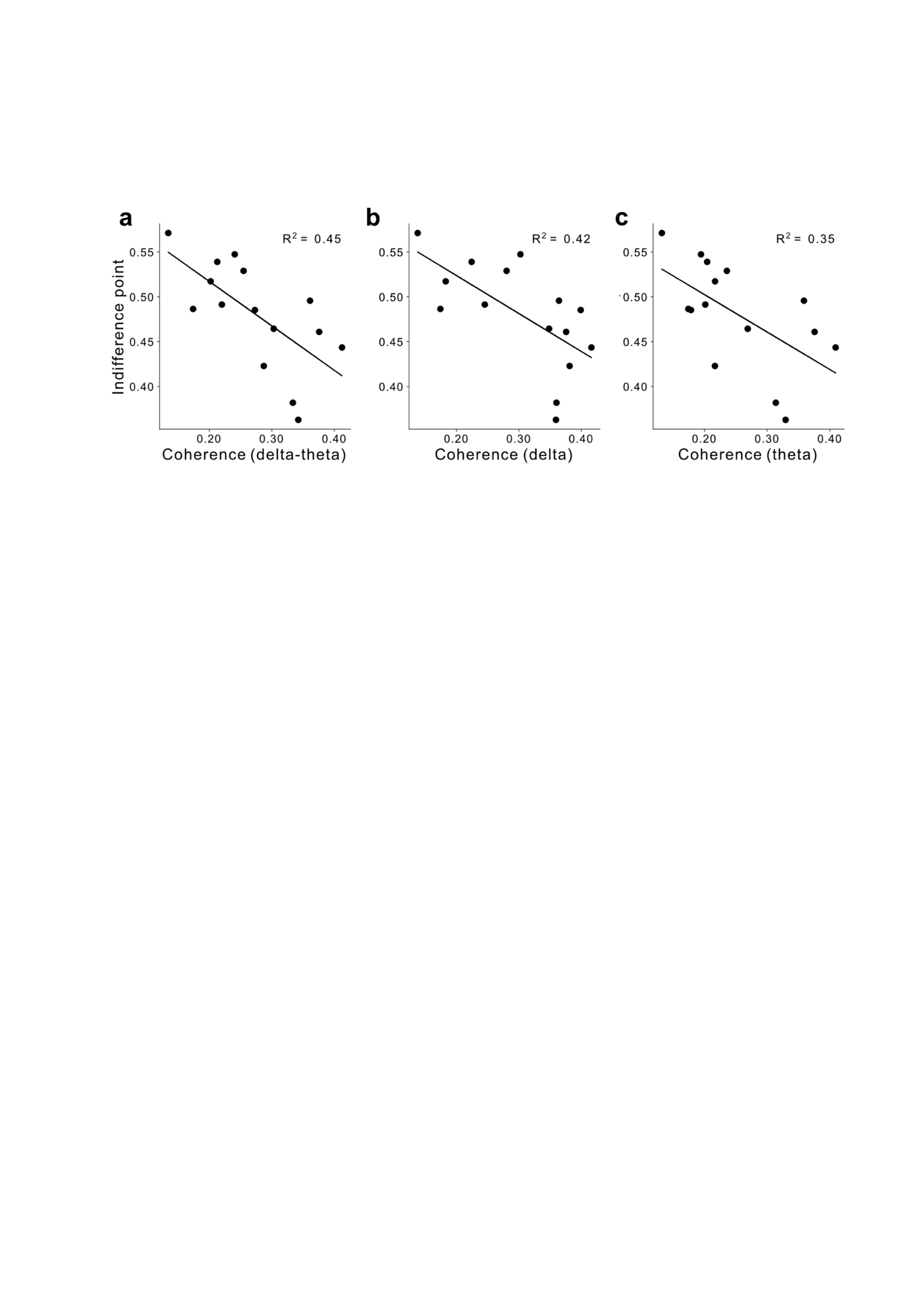

Extended Data Figure 3. Significant correlation between risk preference and OFC coherence in delta-theta, delta, and theta frequency in 3 second time window before subject choice. To ensure that we used a sufficiently long epoch to correctly estimate slow delta oscillations, we extended our analysis window to 3 seconds prior to the button press—encompassing a minimum of three cycles of a waveform in the target frequency. We calculated the mean pairwise coherence among all OFC electrodes for each subject in the delta-theta, delta, and theta frequencies, and performed linear regression with subject’s risk indifference point. a, Delta-theta band (*R^2^* =0.45, *P*=0.0065). b, Delta band (*R^2^* =0.42, *P*=0.009). c, Theta band (*R^2^* =0.35, *P*=0.02).

**Fig. S4.**

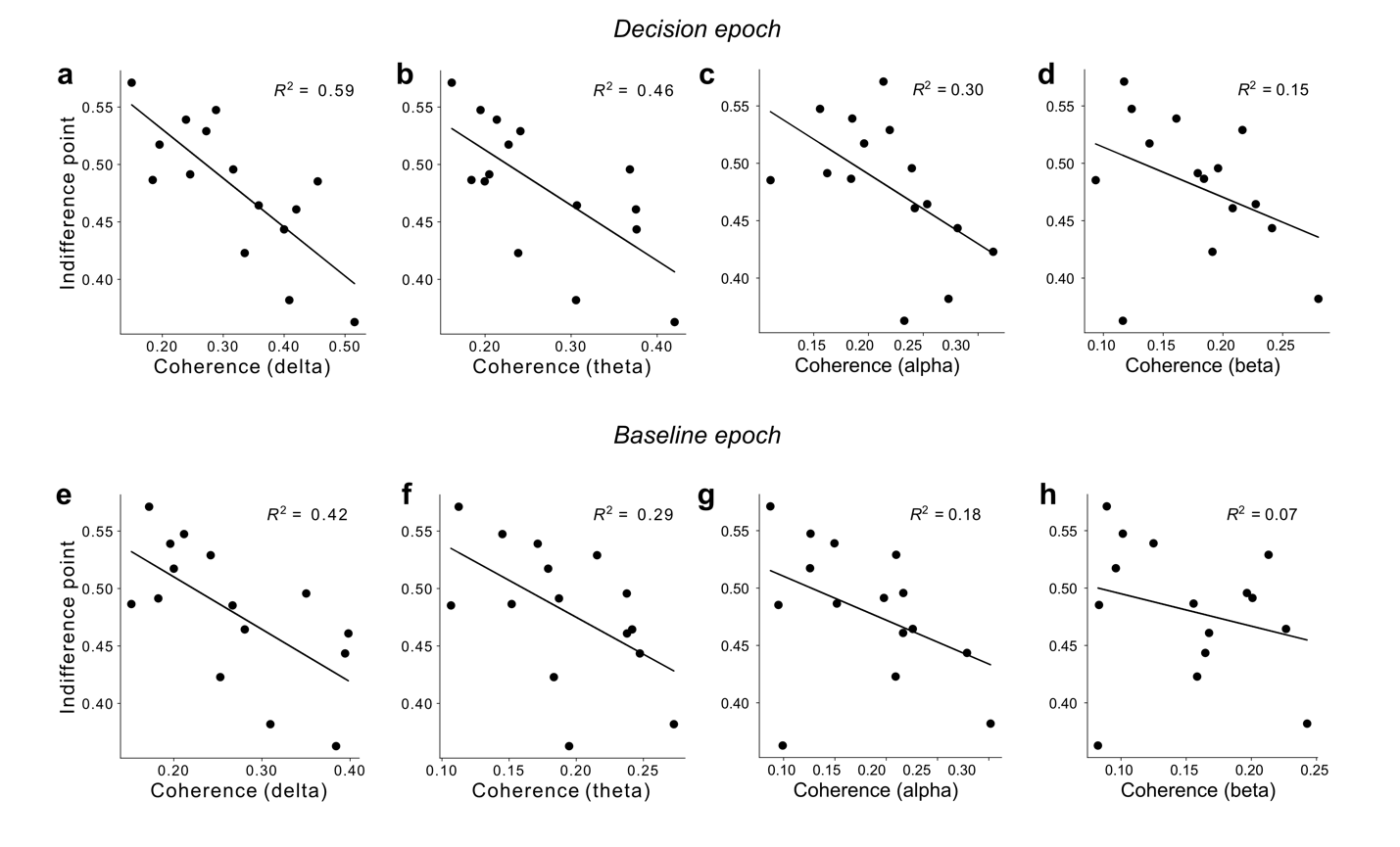

**Extended Data Figure 4**. **Correlation between risk preference and OFC coherence across frequency bands.** We calculated the mean pairwise coherence among all OFC electrodes for each subject in the delta, theta, alpha, and beta frequency separately during the decision epoch and the baseline epoch and performed linear regression with subject’s risk indifference point**. a-d,** Decision epochs: **a**, Significant correlation in delta band (*R^2^* =0.59, *P*=0.0008). **b,** Significant correlation in theta band (*R^2^* =0.46, *P*=0.005). **c,** Weak correlation in alpha band (*R^2^* =0.30, *P*=0.0345). **d,** Non-significant correlation in beta band (*R^2^* =0.15, *P*= 0.152). **e-h,** Baseline epochs: **e,** Significant correlation in delta band (*R^2^* =0.42, *P*=0.009). **f,** Significant correlation in theta band (*R^2^* =0.29, *P*=0.03). **g,** Non-significant correlation in alpha band (*R^2^* =0.18, *P*=0.112). **h,** Non-significant correlation in beta band (*R^2^* =0.07, *P*=0.346).

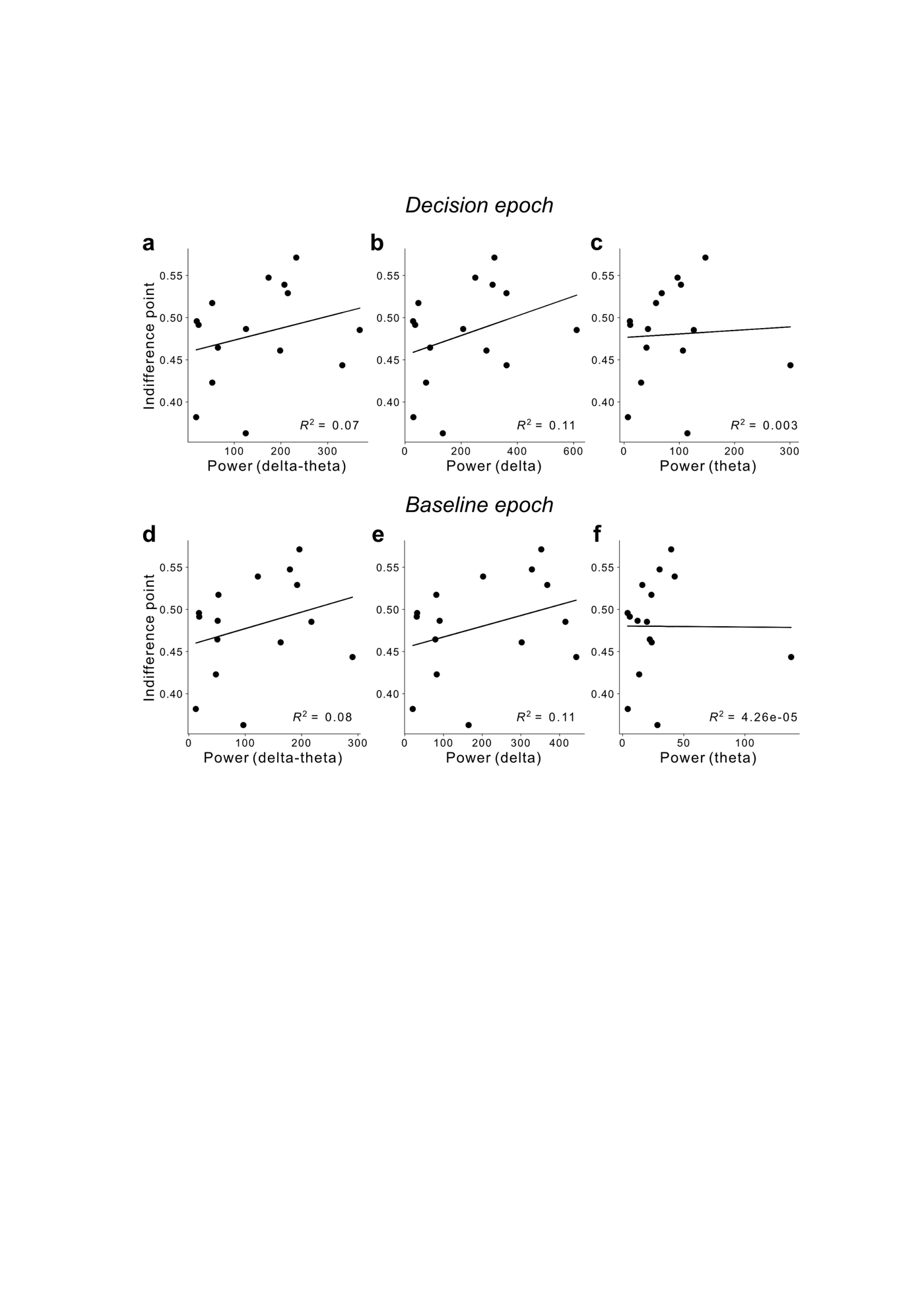
**Fig. S5.**

Extended Data Figure 5. Lack of correlation between risk preference and OFC power in delta-theta, delta, and theta frequencies. To ensure the specificity of this effect to oscillatory phase coherence, rather than mere oscillatory power, we examined the relationship between risk attitudes and power in the delta-theta, delta, and theta frequencies. a, Delta-theta band in decision epoch, *R^2^* =0.07, *P*=0.336. b, Delta band in decision epoch, *R^2^* =0.11, *P*= 0.228. c, Theta band in decision epoch, *R^2^* =0.003, *p*=0.851. d, Delta-theta band in baseline epoch, *R^2^* =0.08, *P*=0.297. e, Delta band in baseline epoch, *R^2^* =0.11, *p*=0.23. f, Theta band in baseline epoch, *R^2^* = 4.26e-05,*P*= 0.982.

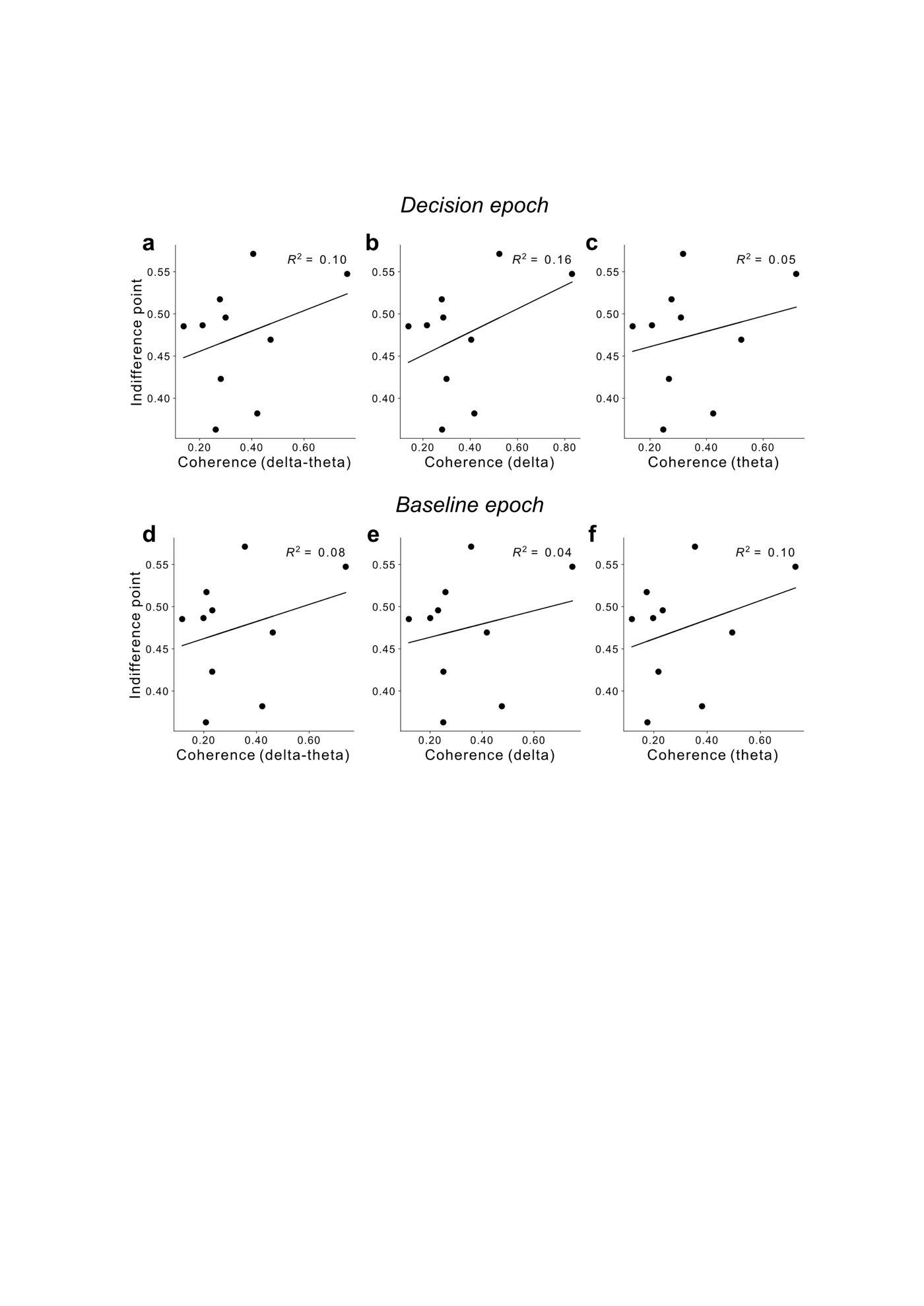
**Fig. S6.**

**Extended Data Figure 6. Lack of correlation between risk preference and hippocampal coherence in delta-theta, delta, and theta frequencies.** To assess the anatomical specificity of our findings, we replicated our analyses in a control region not directly implicated in decision-making under uncertainty, the hippocampus. We calculated the mean pairwise coherence among all hippocampal electrodes for each subject in the delta-theta, delta and theta frequencies during the decision epoch and the baseline epoch, and performed linear regression with subject’s risk indifference point. **a,** Delta-theta band in decision epoch, *R^2^* =0.10, *P*=0.368. **b,** Delta band in decision epoch, *R^2^* =0.16, *P*=0.248. **c,** Theta band in decision epoch, *R^2^* =0.05, *p*=0.519. **d,** Delta-theta band in baseline epoch, *R^2^* =0.08, *P*=0.436. **e,** Delta band in baseline epoch, *R^2^* =0.05, *P*=0.554. **f,** Theta band in baseline epoch, *R^2^* =0.10, *P*=0.366.

**Fig. S7.**

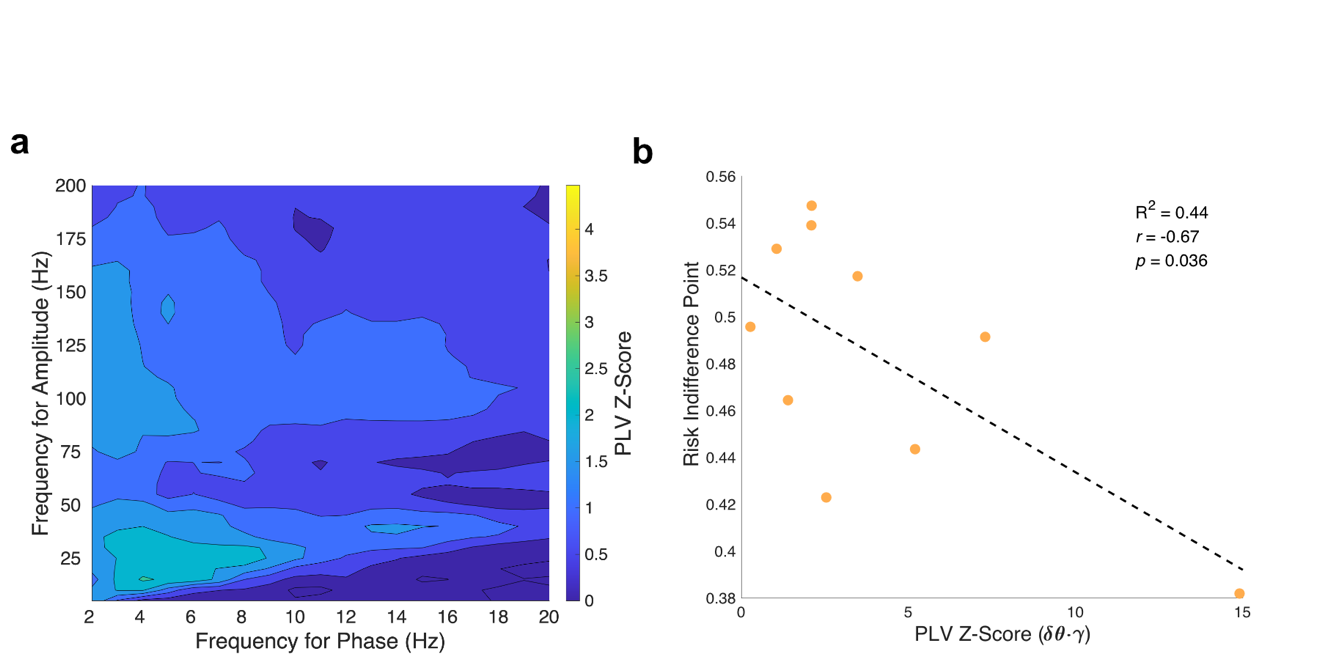

**Extended Data Figure 7. Cross frequency phase-amplitude coupling**

**a,** Group-level comodulogram of mean PLV_z_ from non-significant electrodes across subjects (n=94/130 electrodes across n=15 subjects). Mean PLV_z_ across all phase-amplitude frequency pairs was 1.00 ($\sigma$=0.49, max = 2.57). The peak phase frequency (*lf_p_*) was 4Hz theta and the peak amplitude frequency (*hf_a_*) was 30Hz gamma. **b,** Correlation between risk indifference point and average cross frequency phase-amplitude coupling between delta-theta phase and broadband gamma amplitude in significant electrodes. In the subjects (n=10/15) with electrodes (n=94) showing significant cross frequency phase-amplitude coupling, the subject’s risk indifference point was negatively correlated with PLV_z_ between delta-theta *lf_p_* (δθ, 2-8Hz) and broadband gamma *hf_a_* (Hγ, 30-200Hz; Pearson *r* = -0.67, *p* = 0.036, *R^2^* = 0.444, Adjusted *R^2^* = 0.374).

**Fig. S8.**

**Extended Data Figure 8. White matter stimulation does not alter individual risk preference or reaction times. a,** Anatomical location of electrodes used for stimulation in patient s03. The two deepest contacts were selected for stimulation, and their anatomical location was subsequently verified by examination of the co-registered MR-CT images. The deepest contact was used as the anode in stimulation (red point) but was later identified to be in white matter as was the second deepest contact, used as cathode. **b,** Behavioral risk patterns during gambling task before (black line) and during stimulation (blue line) in patient s03. The plot shows the proportion of gambles that patients chose as a function of the win probability of the gamble offer in 10% increments. The patient showed no difference in risk preference during white-matter stimulation. **c,** Reaction times in patient s03 for baseline (Base, grey points) and stimulation (Stim, blue points) conditions.
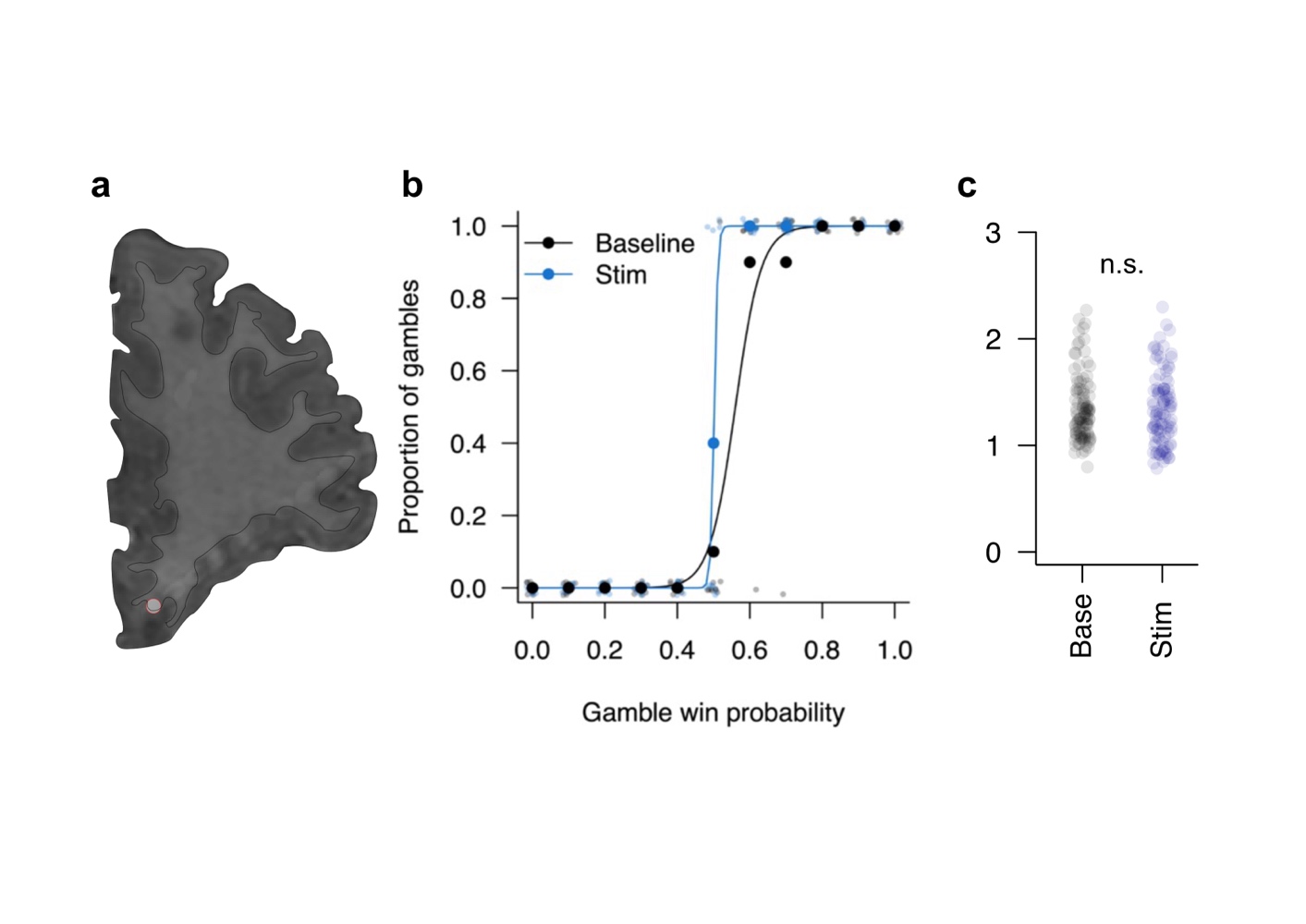

**Table S1.**

| **Subject** | **Age** | **Sex** | **Handedness** | **#OFC Electrodes** |
| --- | --- | --- | --- | --- |
| S01 | 29 | M | R | 5 |
| S02 | 42 | M | R | 49 |
| S03 | 44 | M | R | 7 |
| S04 | 31 | M | L | 11 |
| S05 | 48 | F | - | 10 |
| S06 | 30 | M | R | 14 |
| S07 | 33 | F | R | 3 |
| S08 | 50 | M | R | 5 |
| S09 | 34 | M | R | 3 |
| S10 | 31 | M | L | 4 |
| S11 | 37 | F | L | 3 |
| S12 | 40 | F | L | 4 |
| S13 | 34 | M | A | 2 |
| S14 | 22 | F | R | 6 |
| S15 | 23 | M | R | 4 |

**Extended Data Table 1. Demographic information and number of OFC electrodes in each patient.** M: male; F: female. R: right-handed; L: left-handed; A: ambidextrous.
